# Loss-of-function of ALDH3B2 transdifferentiates human pancreatic duct cells into beta-like cells

**DOI:** 10.1101/2024.05.13.593941

**Authors:** Jian Li, Kevin Bode, Yu-chi Lee, Noelle Morrow, Andy Ma, Siying Wei, Jessica da Silva Pereira, Taylor Stewart, Alexander Lee-Papastavros, Jennifer Hollister-Lock, Brooke Sullivan, Susan Bonner-Weir, Peng Yi

## Abstract

Replenishment of pancreatic beta cells is a key to the cure for diabetes. Beta cells regeneration is achieved predominantly by self-replication especially in rodents, but it was also shown that pancreatic duct cells can transdifferentiate into beta cells. How pancreatic duct cells undergo transdifferentiated and whether we could manipulate the transdifferentiation to replenish beta cell mass is not well understood. Using a genome-wide CRISPR screen, we discovered that loss-of-function of ALDH3B2 is sufficient to transdifferentiate human pancreatic duct cells into functional beta-like cells. The transdifferentiated cells have significant increase in beta cell marker genes expression, secrete insulin in response to glucose, and reduce blood glucose when transplanted into diabetic mice. Our study identifies a novel gene that could potentially be targeted in human pancreatic duct cells to replenish beta cell mass for diabetes therapy.

## INTRODUCTION

Diabetes, no matter the cause or type, is a disease of pancreatic beta cell deficiency ^1^. Current treatments for diabetes do not provide the same degree of exquisite glycemic control as would a sufficient number of functional beta cells, and thus do not prevent the debilitating correlates of long-term diabetes. To cure diabetes, one has to find a way to stop the recurrent autoimmune attack on beta cells (type 1 diabetes) or resolve persistent peripheral insulin resistance (type 2 diabetes), but restoring a sufficient functional beta cell mass is critical to a cure for both types of diabetes. Beta cells can be replenished by transplanting human cadaveric islets or islet-like cells derived from human embryonic stem cells (hESC) or induced pluripotent stem cells (iPSCs) ^2–5^. In most cases, transplanted islet cells are HLA-mismatched with the recipient, and immunosuppression is required to prevent graft rejection, creating other complications such as lack of immune defense against pathogens or tumor formation. Alternatively, it may be possible to promote endogenous pancreatic beta cell regeneration. Promoting the regeneration of a patient’s own beta cells could be a safer strategy for beta cell mass replenishment; for type 1 diabetes autoimmunity would still need to be tamed but there would be no need for immunosuppression for allograft rejection. Beta cell regeneration (reviewed in ^6^) can be achieved by self-duplication ^7^ or transdifferentiation from other pancreatic cell types such as duct cells ^8^, alpha cells ^9,10^ or acinar cells ^11^. Pancreatic beta cell replication is the dominant mechanism of beta cell regeneration in adult rodents ^7^. However in human, the adult beta cell replication rate is extremely low and it was postulated that human beta cells regeneration is achieved mainly by transdifferentiation from pancreatic duct cells ^8^. It has been suggested that pancreatic duct cells may serve as a pool of progenitors for both the islet and acinar tissues after birth and into adulthood ^12–18^. Primary human duct cells expanded in culture and overlaid with Matrigel formed islet buds expressing insulin at mRNA and protein levels ^19^. Although human duct-to-beta cell transdifferentiation has been evidenced by the existence of insulin-expressing cells in the pancreatic duct epithelium, it is a very rare phenomenon that many doubt would be relevant for sufficient beta cell regeneration. However, if its underlying mechanism can be understood, one could potentially manipulate duct-to-beta cell transdifferentiation to a high enough efficiency to replenish functional beta cell mass for the treatment of diabetes. Here we employed a genome-wide CRISPR screen to dissect the mechanism of human duct-to-beta cell transdifferentiation and to identify new therapeutic targets for beta cell mass restoration.

## RESULTS

### A genome-wide CRISPR screen identifies ALDH3B2 as a regulator of human duct-to-beta cell transdifferentiation

Forward genetic screening, the genome-wide CRISPR screening in particular, is a powerful approach to discover novel genes, signaling pathways and the underlying mechanisms of a complex biological phenomenon. Here, we developed a genome-wide CRISPR screening strategy to search for genes that regulate the transdifferentiation of human pancreatic duct cells into beta cells. To ensure efficient gene editing and sufficient cell numbers for the genome-wide screen with sufficient coverage, we chose to use an immortalized pancreatic cell line, PANC-1, a human pancreatic carcinoma cell line of ductal origin that maintains many of the differentiated characteristics of normal mammalian pancreatic ductal epithelium ^20,21^. Our strategy was to use PANC-1 cells solely as a CRISPR screen and target discovery tool, and all the findings from the PANC-1 genetic screen were later validated and characterized in primary human pancreatic duct cells. We reasoned that insulin expression or insulin promoter activation would be the most direct and the simplest readout for cell transdifferentiation into pancreatic beta-like cells. Therefore, we introduced a reporter construct, Rat insulin promoter 3.1 (<u>R</u>IP)-<u>E</u>GFP-<u>P</u>2A-Blasticidin-S deaminase (<u>B</u>SD) (referred to as REPB reporter), into the PANC-1 cells via lentiviral transduction to create a REPB-PANC-1 reporter cell line (Fig. 1a). Rat insulin promoter (RIP) is known to be active in human beta cells and RIP 3.1 is a modified rat insulin promoter that is believed to have higher efficiency and beta cell specificity ^22^, which could increase the sensitivity of our CRISPR screen. The P2A peptide ensures the co-expression of the EGFP and BSD reporter genes, the EGFP reporter allows to visually and quantitatively monitor insulin promoter activation, and the expression of the BSD reporter gene confers resistance to blasticidin treatment, making it easy to enrich or select insulin promoter-activated cells. To validate the REPB reporter construct, we also transduced the NIT-1 mouse beta cell line with the REPB reporter (REPB-NIT-1 cell line). As shown in Extended Data Fig. 1a and 1b, the EGFP expression can only be observed in the REPB-NIT-1 cells, but not at all in REPB-PANC-1 cells. The REPB-PANC-1 cells, but not the REPB-NIT-1 cells, were sensitive to blasticidin treatment (data not shown).

**Fig. 1.**
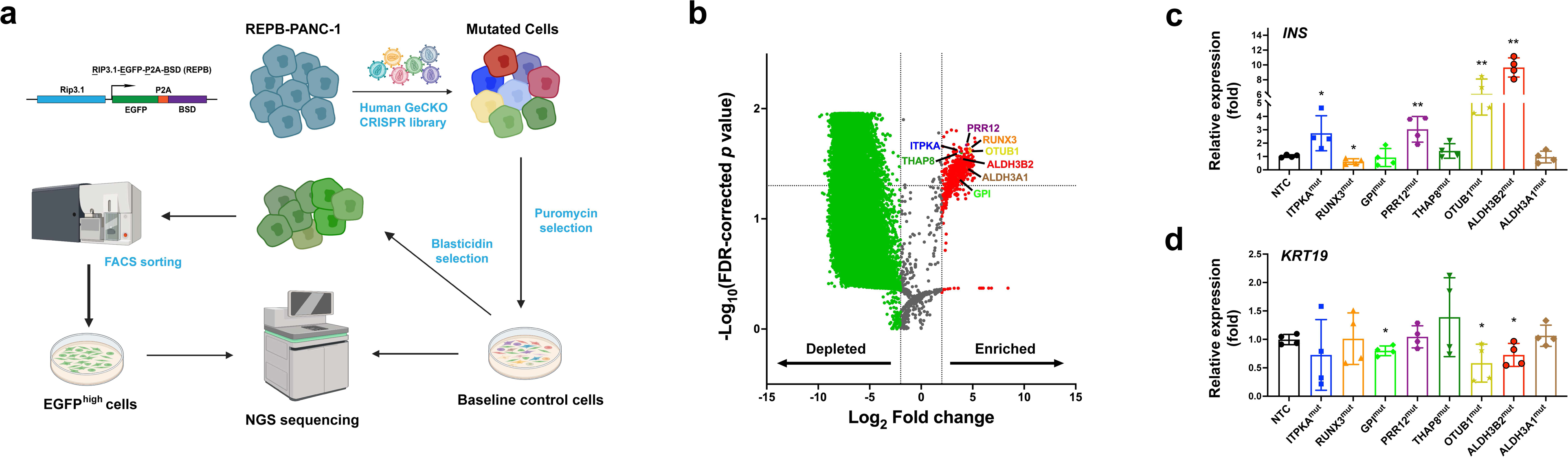
Genome-wide CRISPR screen identifies ALDH3B2 as a regulator of pancreatic duct-to-beta cell transdifferentiation. **a,** The structure of REPB reporter and an illustration of genome-wide CRISPR screen workflow. The REPB reporter contains Rat insulin promoter (RIP3.1) driving the expression of EGFP and Blasticidin-S deaminase (BSD), the EGFP and BSD genes are fused together with P2A peptide. **b,** gRNA profile volcano plots of the genome-wide CRISPR screen showing the depleted and enriched gRNA in the EGFP^high^, blasticidin resistance REPB-PANC-1 cells. The x axis is the fold change of gRNA counts (read per million, CPM), and the y axis is statistical significance as shown by -log_10_ of the false discovery rate corrected p value. The vertical dashed line represents a fold change for gene threshold of 1, the horizontal dashed line represents a p value threshold of 0.05. Several highly enriched gRNAs are labeled as large dots with various colors; up-regulated gRNAs were shown as small red dots and down-regulated gRNAs as small green dots; all unchanged gRNAs were as small gray dots. **c and d:** qPCR analysis of INS **(c)** and KRT19 **(d)** expression level for top 8 candidate gene mutant PANC-1 cells.

To execute the CRISPR screen, as illustrated in Fig. 1a, we transduced the REPB-PANC-1 cells with a human lentiviral genome-wide CRISPR knockout library (GeCKO v2) ^23^ that comprises approximately 120,000 guide RNAs (gRNAs) targeting a total of 19,050 genes. We used a low multiplicity of infection (MOI) ∼0.3 to ensure that most of the cells carry only one mutation. Briefly, approximately 10^8^ lentiviral library transduced REPB-PANC-1 cells were treated with low dose of blasticidin (10ug/ml) for 7 days, and then the blasticidin resistant cells were subjected to FACS sorting based on their EGFP intensity (Extended Data Fig. 2). Using next generation sequencing (NGS) and bioinformatic analysis, the gRNA profile of the highest EGFP-expressing cells (EGFP ^high^, blasticidin-resistant) was generated and compared to that of the cells without blasticidin selection and FACS sorting (Supplemental Table 1). As shown in Fig. 1b, the volcano plot showed that the majority of the gRNAs were depleted from the EGFP^high^, blasticidin-resistant REPB-PANC-1 cells, whereas only small number of gRNAs were significantly enriched.

To narrow down the candidate gRNAs for validation, we selected all the gRNAs that are significantly enriched in the EGFP^high^ cells (FDR<0.05), and then ranked them based on their <u>c</u>ount <u>p</u>er <u>m</u>illion (CPM). We then picked the top 8 candidate gRNAs from the list and generated individual mutant PANC-1 cell lines using the corresponding gRNAs identified in our screen. We used quantitative PCR (qPCR) to analyze the expression of endocrine marker genes including insulin (INS), glucagon (GCG) and somatostatin (SST), as well as pancreatic duct cell marker gene keratin 19 (KRT19 or CK19). Several mutant PANC-1 cell lines showed differential expression of the examined marker genes (Fig. 1c, 1d and Extended Data Fig. 3a and 3b). In particular, PANC-1 cells transduced with a gRNA targeting ALDH3B2 showed the highest INS expression and the lowest KRT19 levels compared to non-targeting control (NTC) gRNA transduced PANC-1 cells. ALDH3B2, also known as ALDH8, is one of 19 members of the human aldehyde dehydrogenase (ALDH) superfamily that convert various types of aldehydes to carboxylic acids ^24^. It is well documented that ALDH genes are important regulators of stem cells and cell fate determination ^25^. Another close member in the ALDH family, ALDH1A3, was recently shown to contribute to pancreatic beta cell failure and de-differentiation in type 2 diabetes ^26^. We reasoned that ALDH3B2, as an enzyme, could potentially be an easier therapeutic target for small molecules targeting. Therefore, we prioritized ALDH3B2 for further in-depth validation and characterization.

### Loss-of-function of ALDH3B2 trans-differentiates PANC-1 cells into pancreatic beta-like cells

We generated an ALDH3B2 mutant PANC-1 cell line by lentiviral transduction of SpCas9 and ALDH3B2 gRNA into PANC-1 cells. Genomic sequencing of the targeted region in ALDH3B2^mut^ PANC-1 cells revealed that more than 75% of the sequenced ALDH3B2 alleles carried indel mutations (Extended Data Fig. 4). Western blot confirmed that ALDH3B2 protein level in the ALDH3B2^mut^ PANC-1 cells was reduced by ∼40% compared to the non-targeting-control (NTC) gRNA lentivirus transduced PANC-1 cells (Extended Data Fig. 5a and 5b). To ensure that loss-of-function of ALDH3B2 induces *bona fide* cell transdifferentiation and did not only just activate the insulin promoter, we conducted a series of qPCR experiments to examine additional genes characteristic for pancreatic beta cells. We found that the expression of pancreatic endocrine hormones, insulin (INS) and somatostatin (SST), but not glucagon (GCG) or pancreatic polypeptide (PP), were significantly increased in the ALDH3B2 mutant PANC-1 cells (Fig. 2a). In addition, the expression of several of key beta cell transcription factors, including *PDX1, MAFA, NGN3* and *PAX6*, were also significantly upregulated in the ALDH3B2 mutant PANC-1 cells (Fig. 2b). We examined additional genes that are critical for beta cell function and found that *GLUT1 (SLC2A1), GLUT2 (SLC2A2), GLUCOKINASE (GCK)*, subunits of the ATP-sensitive potassium (K-ATP) *(KCNJ11* and *ABCC8*), *CPE* and *IA2* were all significantly upregulated in the ALDH3B2 mutant PANC-1 cells (Fig. 2c). ALDH3B2 mutant PANC-1 cells had slightly decreased expression of pancreatic duct markers KRT19 and CA2 but not of HNF1B or Sox9 (Fig. 2d). Immunofluorescence imaging showed that clusters of ALDH3B2 mutant PANC-1 cells expressed human Insulin (INS) and C-peptide (CPEP), whereas neither insulin and C-peptide were detected in control PANC-1 cells (Fig. 2e). The insulin content of the ALDH3B2 mutant PANC-1 was significantly increased compared to control cells (Fig. 2f). In addition, using electron microscopy (EM), we found that many of the ALDH3B2 mutant PANC-1 cells had insulin granules (vesicles with halo, arrows in Fig. 2g), while no insulin granules were detectable in control -PANC-1 cells (Fig. 2g). Intriguingly, the ALDH3B2 mutant PANC-1 cells had intensive endoplasmic reticulum (ER) network (Fig. 2g), a characteristic found in pancreatic beta cells but not pancreatic duct cells. Collectively, these studies indicate that loss-of-function mutations in ALDH3B2 in PANC-1 cells not only trigger insulin promoter activation but also precipitate a significant cell fate transformation, shifting from a pancreatic ductal phenotype to a beta-like profile.

**Fig. 2.**
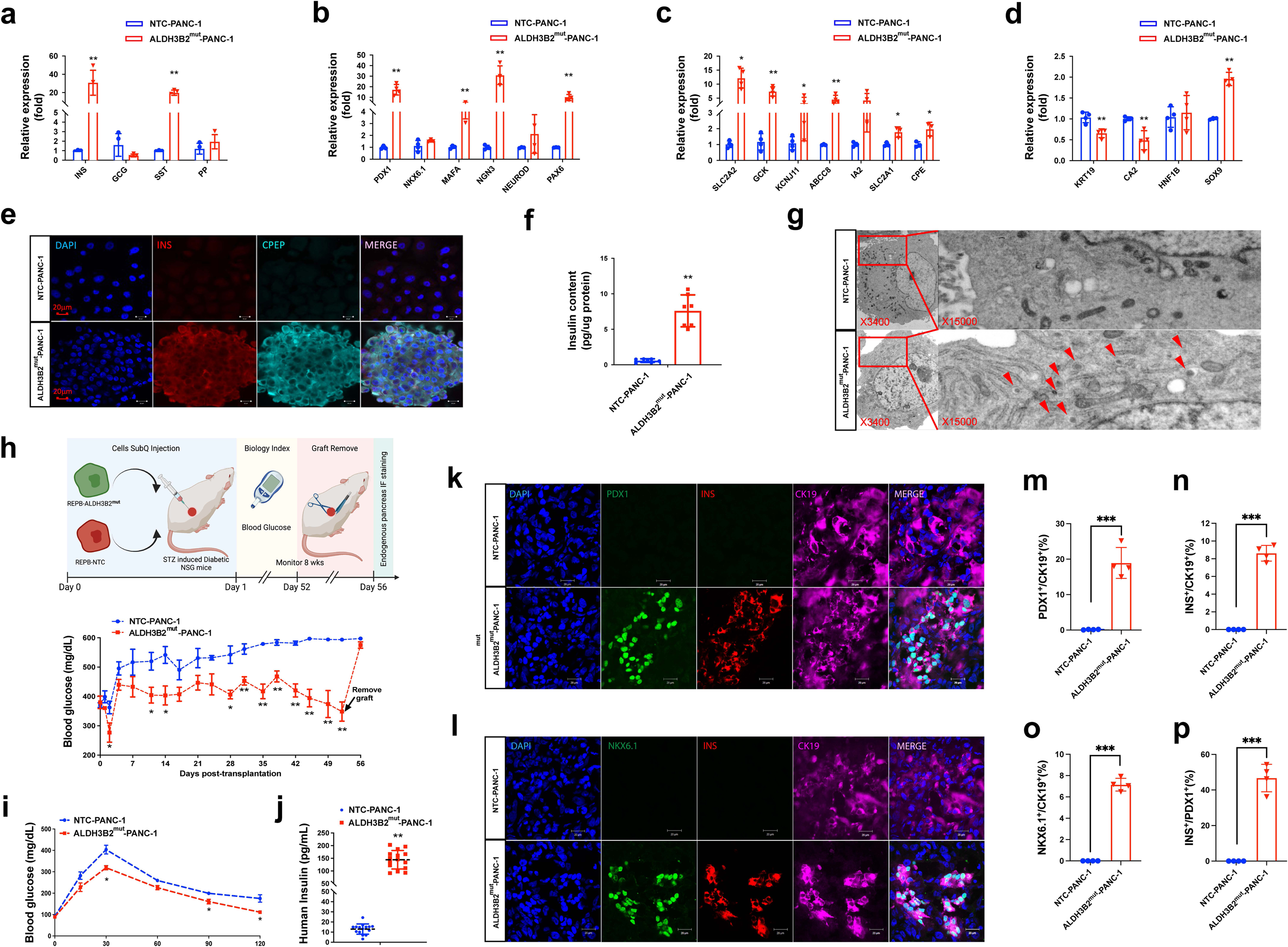
Loss-of-function of ALDH3B2 trans-differentiates PANC-1 cells into insulin-producing beta-like cells. **a-d:** qPCR analysis of human beta cell and duct signature genes. Data show mean ± SEM of n=4 biological replicates per condition and are representative of 2–3 independent experiments. *p < 0.05, **p < 0.01, calculated by two-way ANOVA with Sidak’s multiple comparisons test. **a,** Genes related to pancreatic endocrine hormones. **b,** Genes related to beta cell key transcription factors. **c,** Genes related to beta cell function genes. **d,** Genes related to pancreatic duct cell markers. **e,** Insulin and C-peptide immunofluorescence in NTC cells and ALDH3B2^mut^-PANC-1 cells. **f:** Insulin content in NTC cells and ALDH3B2^mut^-PANC-1 cells were measured by insulin ELISA, normalized by cells genomic DNA content. n=7 biological replicates per condition and genotype. Data show mean ± SEM, **p < 0.01. **g**, Electron microscopy (EM) analysis of NTC-PANC-1 cells and ALDH3B2^mut^-PANC-1 cells. Insulin secretion vesicles are indicated by arrow heads. **h**, An illustration of the in vivo function evaluation of trans-differentiated PANC-1 cells, and random blood glucose levels in the NTC and ALDH3B2^mut^-PANC-1 cells transplanted diabetic NSG mice over 8 weeks of monitoring. Data show mean ± SEM of n=5 mice (NTC-PANC-1) and n=6 mice (ALDH3B2^mut^-PANC-1) per group and are representative of three experiments, calculated by two-way ANOVA with Tukey’s multiple comparisons test. **i,** An Intraperitoneal glucose tolerance test (IPGTT) in NTC or ALDH3B2^mut^-PANC-1 cells transplanted NSG mice 6 weeks post-transplantation. **j,** Serum human insulin levels measurement in the NTC and ALDH3B2^mut^-PANC-1 cells transplanted diabetic NSG mice. **k,** Pdx1 and Insulin immunofluorescence in transplanted NTC cells and ALDH3B2^mut^-PANC-1 cells. **l,** Nkx6.1 and Insulin immunofluorescence in transplanted NTC cells and ALDH3B2^mut^-PANC-1 cells. **m-o,** Percentage of Pdx1^+^ **(m)**, Insulin^+^ **(n)**, Nkx6.1^+^ **(o)** cells in all CK19^+^ cells. **p,** Percentage of Insulin^+^ cells in all Pdx1^+^ cells.

To evaluate whether these trans-differentiated pancreatic beta-like cells were functional, we performed *in vitro* <u>g</u>lucose <u>s</u>timulated insulin <u>s</u>ecretion (GSIS) assay comparing control NTC-PANC-1 cells and transdifferentiated ALDH3B2 mutant PANC-1 cells, and we found that the ALDH3B2 mutant PANC-1 cells can secrete significantly more insulin at baseline (2.8 mM glucose) compared to the control NTC-PANC-1 cells, and also mildly respond to higher glucose (16.7 mM glucose) (Extended Data Fig. 6a). To test whether the ALDH3B2 mutant PANC-1 cells were also functional *in vivo*, where and then we transplanted the ALDH3B2 mutant or control PANC-1 cells (NTC) subcutaneously into STZ diabetic mice (Fig. 2h, upper panel). Mice transplanted with the ALDH3B2 mutant PANC-1 cells showed significantly decreased daily random blood glucose (Fig. 2h, lower panel) and improved glucose tolerance (Fig. 2i) compared to mice transplanted with NTC-PANC-1 cells. Notably, when the ALDH3B2 mutant PANC-1 graft was removed at the end of the study, blood glucose increased to the same level as in the control mice, confirming that the blood-glucose-lowering was indeed caused by the transplanted ALDH3B2 mutant PANC-1 cells (Fig. 2h, lower panel). Human insulin serum levels were also significantly higher in mice transplanted with ALDH3B2 mutant PANC-1 cells (Fig. 2j). Immunofluorescent imaging showed that transplanted ALDH3B2 mutant PANC-1 cells co-expressed PDX1, Insulin, NKX6.1, C-peptide and CK19 (Fig. 2k, 2l, Extended Data Fig. 7a and 7b). We observed that a few cells co-express somatostatin (SST) and Insulin, but no cell expresses glucagon (GCG), or the exocrine cell marker gene amylase (AMY) (Extended Data Fig. 7c and 7d). Of all the transplanted ALDH3B2 mutant PANC-1 cells, ∼20% were PDX1^+^ (Fig. 2m) and ∼8% were INS^+^ or NKX6.1^+^ (Fig. 2n and 2o). Almost all the INS^+^ cells were also NKX6.1^+^, suggesting that trans-differentiated beta-like cells adopted a true beta cell phenotype. It should also be noted that only ∼45% of the PDX1^+^ cells co-expressed insulin (Fig. 2p), and we speculate that the PDX1^+^/INS^-^ cells may represent pancreatic progenitor-like cells that have yet committed to beta cell fate.

We employed an inducible shRNA system to ensure that the transdifferentiation of PANC-1 cells into beta-like cells by ALDH3B2 CRISPR knockout was indeed due to the loss-of-function of ALDH3B2 and not caused by off-target effects of the ALDH3B2 gRNA. We generated PANC-1 cell lines carrying a Tet-On inducible ALDH3B2 shRNA or a scrambled control shRNA (Fig. 3a, left panel). The ALDH3B2 shRNA PANC-1 cells with Doxycycline treatment showed significantly reduced ALDH3B2 mRNA expression (Fig. 3a right panel) and protein level (Extended Data Fig. 5c and 5d) after doxycycline (dox) treatment. Similar to the ALDH3B2 CRISPR mutant PANC-1 cells, knock-down of ALDH3B2 by shRNA also trans-differentiated PANC-1 cells into beta-like cells. A series of qPCR experiments showed that the expression of key beta cell transcription factors including *PDX1*, *MAFA, NGN3, NEUROD* and *PAX6* (Fig. 3b), endocrine hormone insulin (*INS*) and somatostatin (*SST*) (Fig. 3C), and beta cell function related genes (*SLC2A2, GCK, KCNJ11* and *ABCC8*) were significantly increased (Fig. 3d), whereas the expression of pancreatic duct cell marker genes (*KRT19, CA2* and *SOX9*) were reduced (Fig. 3e). Human insulin could also be detected in shALDH3B2 PANC-1 cells (+Dox) but not in shControl PANC-1 cells or in shALDH3B2 PANC-1 cells (+Dox) by immunofluorescence (Fig. 3f). These analyses confirmed that loss-of-function of ALDH3B2 by CRISPR targeting or shRNA silencing allowed PANC-1 cells to transdifferentiate and adopt a beta-like cell fate.

**Fig. 3.**
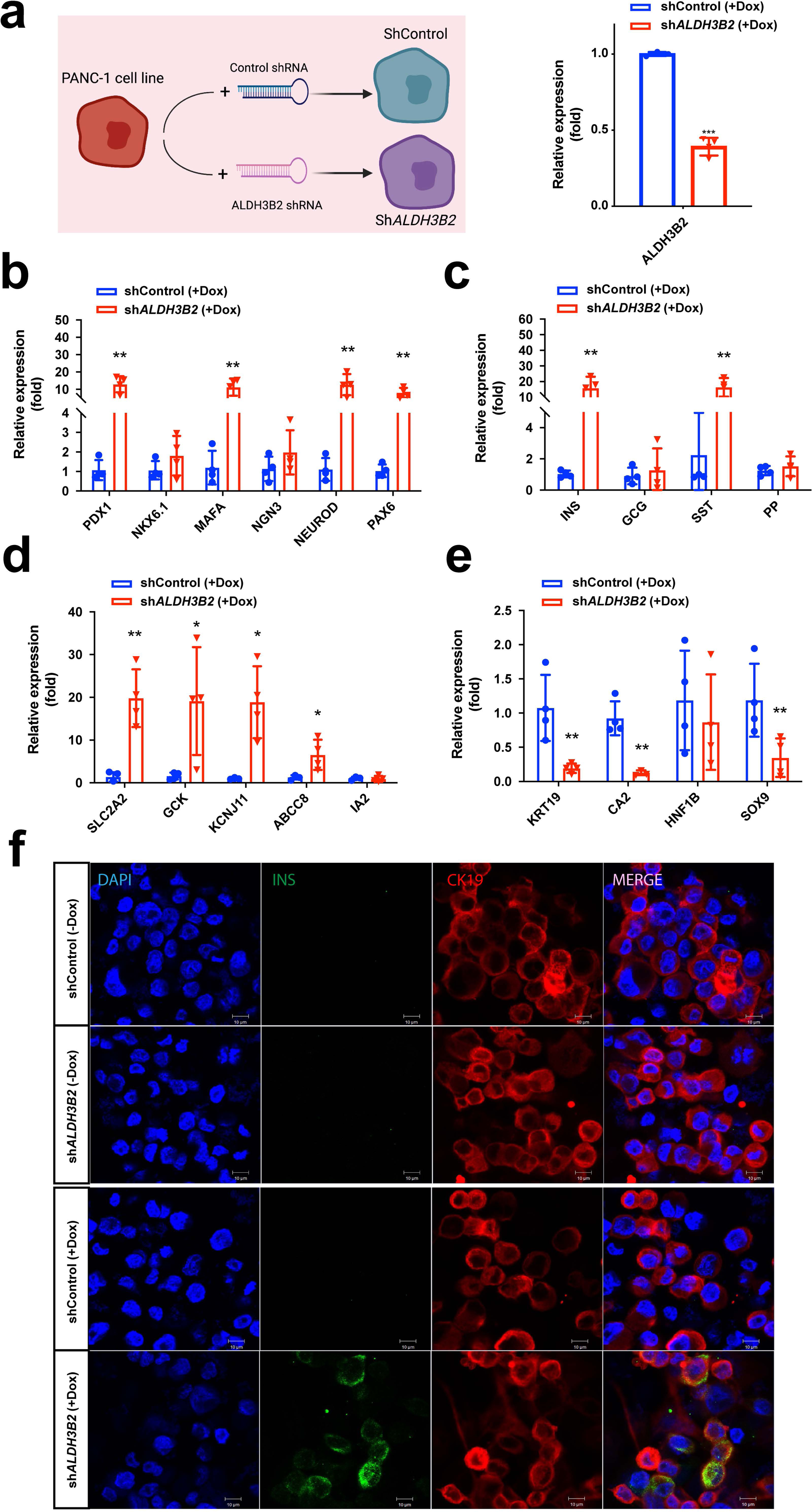
Knock-down of ALDH3B2 by shRNA trans-differentiates PANC-1 cells into beta-like cells. **a,** Generation of inducible non-targeting (shControl) cells and shALDH3B2-PANC-1 cells, and ALDH3B2 expression is significantly reduced in shALDH3B2 PANC-1 cells with Doxycycline treatment shown by qPCR analysis. **b-e:** qPCR analysis of human beta cell and duct signature genes. Data show mean ± SEM of n=4 biological replicates per condition and are representative of 2–3 independent experiments. *p < 0.05, **p < 0.01, calculated by two-way ANOVA with Sidak’s multiple comparisons test. **b,** Genes related to beta cell key transcription factors. **c,** Genes related to pancreatic endocrine hormones. **d,** Genes related to beta cell function genes. **e,** Genes related to pancreatic duct cell markers. **f,** Insulin and C-peptide immunofluorescence imaging in shControl and shALDH3B2 PANC-1 cells with or without Doxycycline treatment.

### Loss-of-function of ALDH3B2 transdifferentiates human primary pancreatic duct cells into beta-like cells

Next, we tested whether loss-of-function of ALDH3B2 was also able to transdifferentiate <u>h</u>uman <u>p</u>rimary <u>p</u>ancreatic <u>d</u>uct (HPPD) cells into beta-like cells. HPPD cells were isolated and affinity-purified from human donor islet-depleted pancreatic exocrine (listed as “acinar tissue”) from Integrated Islet Distribution Program (IIDP) ^27^. qPCR analyses confirmed lack of insulin expression (Fig. 4a) and high expression of the pancreatic duct marker *KRT19* (Fig. 4b) in the purified HPPD cells compared to primary human islets. Interestingly, we observed that ALDH3B2 expression levels were markedly lower in human islets compared to pancreatic duct cells (Fig. 4c). This differential expression pattern aligns with our results where the mutation of ALDH3B2 in human pancreatic duct cells promotes their transdifferentiation into beta-like cells. These findings suggest that the reduction of ALDH3B2 could be a critical step in the cellular reprogramming process leading to a beta-cell phenotype.

**Fig. 4.**
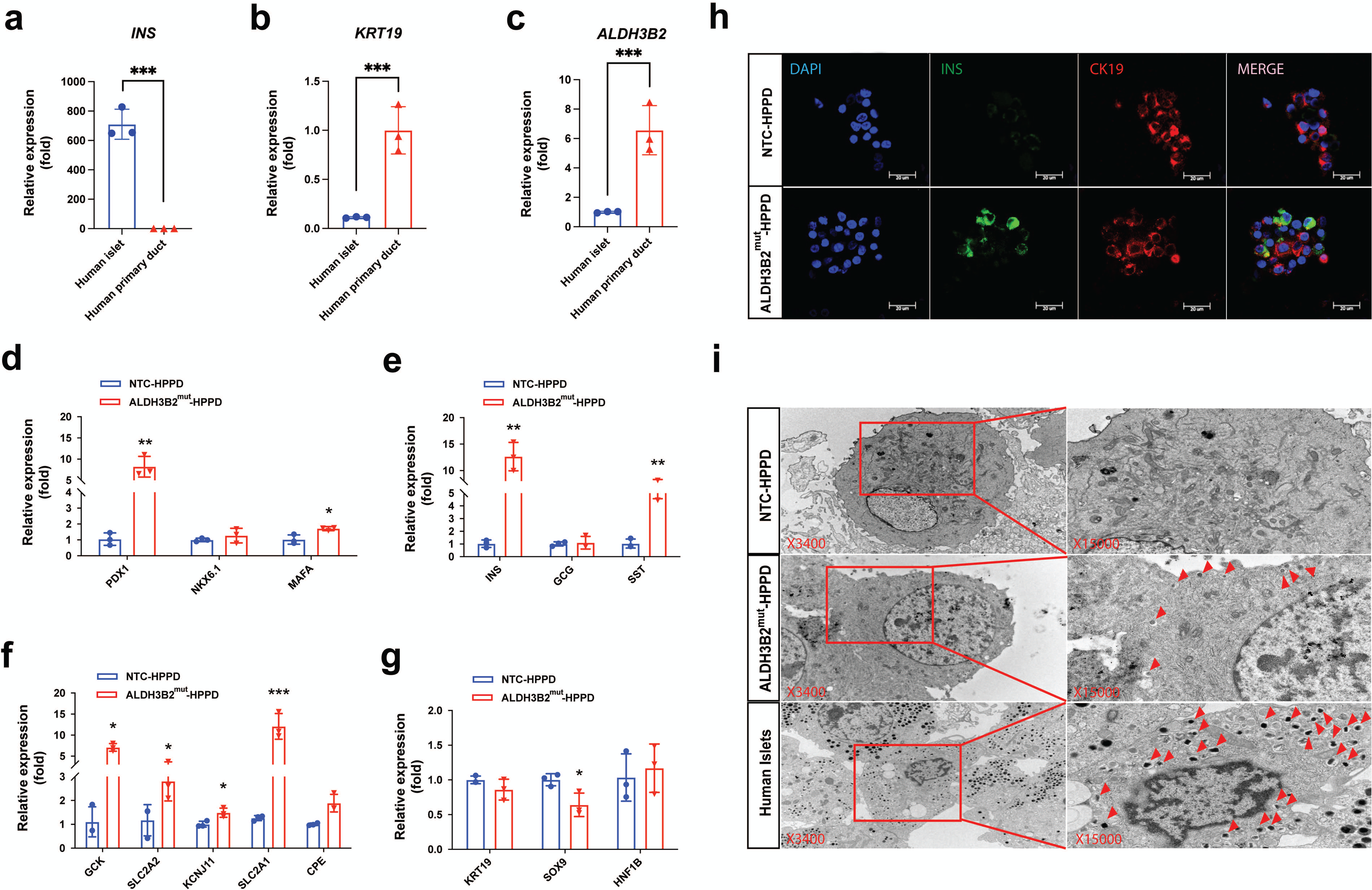
Loss-of-function of ALDH3B2 trans-differentiates human primary pancreatic ductal cells into beta-like cells. **a-c:** qPCR analysis of human Insulin **(a)**, KRT19 **(b)** and ALDH3B2 **(c)** expression in purified human primary pancreatic duct cells (HPPD) and human primary islets. Data show mean ± SEM of n=3. ***p < 0.005. **d-g**, qPCR analysis of human beta cell and duct signature genes. Data show mean ± SEM of n=3 biological replicates per condition and are representative of 2 independent experiments. *P < 0.05, **P < 0.01, calculated by two-way ANOVA with Sidak’s multiple comparisons test. **d,** Genes related to beta cell key transcription factors. **e,** Genes related to pancreatic endocrine hormones. **f,** Genes related to beta cell function genes. **g,** Genes related to pancreatic duct cell markers. **h,** Insulin and CK19 immunofluorescence in NTC-HPPD and ALDH3B2^mut^-HPPD cells. **i**, Electron microscopy (EM) analysis of NTC-HPPD cells, ALDH3B2^mut^-HPPD cells and human primary islet cells. Arrow heads points to insulin secretion vesicles.

Purified HPPD cells were transduced with lentiviruses carrying SpCas9 and either ALDH3B2 gRNA (ALDH3B2^mut^ -HPPD) or a non-targeting control gRNA (NTC-HPPD). qPCR analysis comparing these transduced duct cells showed ALDH3B2^mut^ -HPPD cells had significantly higher expression of key beta cell transcription factors (*PDX1* and *MAFA*) (Fig. 4d), endocrine hormone insulin (*INS*) and somatostatin (*SST*) (Fig. 4e) and beta cell function-related genes (*GCK, SLC2A1, SLC2A2, KCNJ11* and *CPE*) (Fig. 4f). The expression of several pancreatic duct cell marker genes was either unchanged (*KRT19* and *HNF1B*) or slightly reduced (*SOX9*) (Fig. 4g). Using immunofluorescent imaging, we found that a small fraction (∼10%) of HPPD-ALDH3B2^mut^ cells expressed Insulin while still retaining CK19 expression, a possible signature of newly transdifferentiated beta cells from pancreatic duct cells, whereas no insulin expression could be detected in HPPD-NTC cells (Fig. 4h). Furthermore, we also examined whether the ALDH3B2^mut^ -HPPD cells have insulin granules using electron microscopy (EM). Although not as many insulin granules as in primary human beta cells, the ALDH3B2^mut^ -HPPD cells do have significant amount of mature insulin granules, while no insulin granules were detectable in the control NTC-HPPD cells (Fig. 4i).

We performed *in vitro* GSIS assay to evaluate the function of the ALDH3B2^mut^ -HPPD cell and found that compared to control NTC-HPPD cells, the ALDH3B2^mut^ -HPPD cells secreted significantly more insulin and mildly responded to high glucose (Extended Data Fig. 6b). The level of insulin secretion from the ALDH3B2^mut^ -HPPD cells was much lower than human islets (Extended Data Fig. 6b), which we suspect was due to both the relatively low efficacy of the transdifferentiation and the immaturity of the *in vitro* trandifferentiated beta-like cells. We then transplanted ALDH3B2^mut^ -HPPD or NTC-HPPD cells under the kidney capsule of STZ induced diabetic NSG mice and monitored their blood glucose over time (Fig. 5a, upper panel). Within one week mice transplanted with HPPD-ALDH3B2^mut^ cells had significantly lower blood glucose than those mice transplanted with NTC-HPPD cells (Fig. 5a, lower panel). When the ALDH3B2 mutant HPPD grafts were removed at 56 days post-transplantation, blood glucose levels increased to those of the control HPPD transplanted mice, suggesting that the blood-glucose-lowering effect was indeed conferred by the transplanted ALDH3B2 mutant HPPD cells (Fig. 5a, lower panel). Importantly, transplanted ALDH3B2^mut^ -HPPD cells secreted human insulin in response to glucose challenge. We detected a significantly increase in serum human insulin 5 minutes after glucose injection both at 1 week and 3 weeks post-transplantation. No such response was observed in mice transplanted with NTC-HPPD cells or in non-transplanted control NSG mice (Fig. 5b and 5c).

**Fig. 5.**
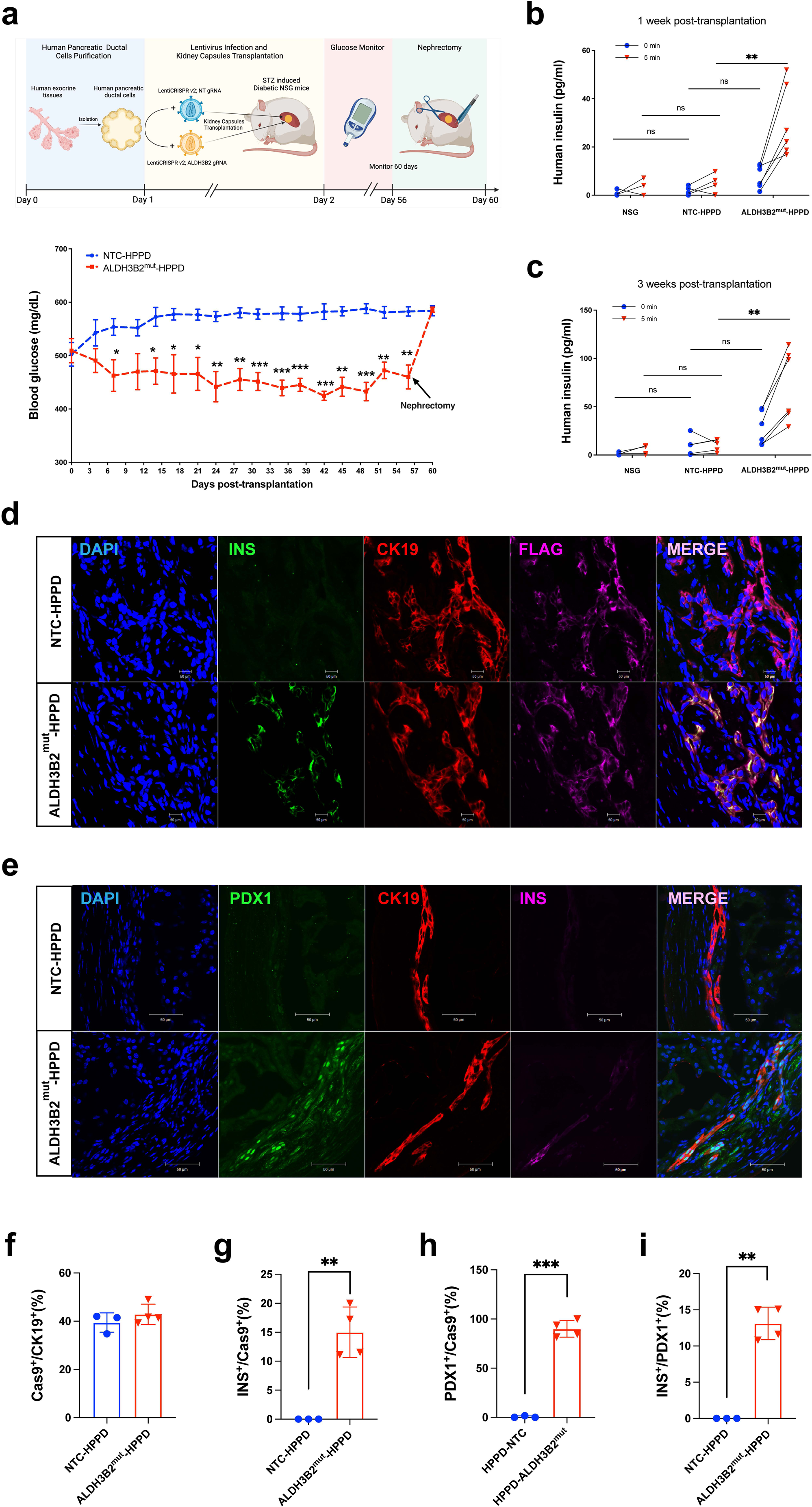
ALDH3B2 mutation trans-differentiated human primary pancreatic duct cells are functional *in vivo*. **a,** An illustration of the in vivo function evaluation of NTC-HPPD cells and ALDH3B2^mut^-HPPD cells, and random blood glucose levels in the NTC-HPPD cells and ALDH3B2^mut^-HPPD cells transplanted diabetic NSG mice for 60 days of monitoring. Data show mean ± SEM of n=5 mice per group, calculated by two-way ANOVA with Tukey’s multiple comparisons test. **b and c,** Serum human insulin levels in non-transplanted NSG mice (Normal NSG), NTC-HPPD cells and ALDH3B2^mut^-HPPD cells transplanted NSG mice 5 minutes after intraperitoneal (IP) glucose injection at 1 week post-transplantation **(b)** and **3** weeks post-transplantation **(c).** Calculated by two-way ANOVA with Sidak’s multiple comparisons test. * NSG vs. NTC-HPPD; # NSG vs. ALDH3B2^mut^-HPPD. **d**, Insulin, CK19 and Cas9 (Flag) immunofluorescence of transplanted NTC-HPPD and ALDH3B2^mut^-HPPD cells. **e**, Pdx1, CK19 and Cas9 (Flag) immunofluorescence of transplanted NTC-HPPD and ALDH3B2^mut^-HPPD cells. **f-i**, percentage of Cas9 lentivirus transduced cells in all transplanted duct cells **(f)**, Insulin^+^ cells in all Cas9 lentivirus transduced cells **(g)**, Pdx1^+^ cells in all Cas9 lentivirus transduced cells **(h)** and Insulin^+^ cells in all Pdx1^+^ cells **(i)**.

While the lack of normalization of glycemia in these experiments may be due to transdifferentiated beta-like cells having a different glucose sensitivity or set-point than true pancreatic beta cells, it is more likely that the number of fully transdifferentiated cells transplanted was inadequate to completely normalized the blood glucose. The issue of higher glucose set-point than primary pancreatic beta cells seems unlikely since when ALDH3B2^mut^-HPPD cells were transplanted into euglycemic NSG mice they improved the glucose tolerance (Extended Data Fig. 8a) and secrete human insulin in response to glucose challenge (Extended Data Fig. 8b and 8c).

We addressed the question of what became of the transplanted cells by immunofluorescent analysis. The grafts had Insulin^+^ cells, C-Peptide^+^ cells, PDX1^+^ cells, NKX6.1^+^ cells only in mice transplanted with ALDH3B2^mut^ -HPPD cells and not with NTC-HPPD cells (Fig. 5d, 5e, Extended Data Fig. 9a-d). Approximately 40% of the transplanted cells were successfully transduced with the NTC or ALDH3B2 gRNA lentivirus as shown by quantification of the percentage of Cas9 (Flag-tagged)^+^/CK19^+^ cells, Fig. 5d and 5f), and almost all of the transplanted ALDH3B2^mut^ -HPPD cells expressed pancreatic duct marker genes CK19 and SOX9 (Extended Data Fig. 9e and 9f). Of those cells infected, ∼15% of ALDH3B2^mut^ - HPPD cells expressed insulin (Fig. 5g) but few, if any, expressed glucagon (GCG), Somatostatin (SST) or Amylase (AMY) (Extended Data Fig. 9e and 9f) . Interestingly, we found that the majority of the pancreatic duct cells infected with ALDH3B2 gRNA lentivirus co-expressed PDX1 and CK19 (Fig. 5e and 5h), and approximately 12% of the PDX1^+^ cells co-expressed insulin (Fig. 5i). Co-expression of PDX1 and CK19 is a signature of pancreatic progenitor cells ^28^, and we postulate that ALDH3B2 loss-of-function may cause the de-differentiation of mature duct cells into pancreatic progenitor-like cells, a portion of which then subsequently differentiate into beta-like cells.

### Loss of ALDH3B2 function in pancreatic duct cells causes epigenetic changes

We found that ALDH3B2 loss-of-function allowed pancreatic duct cells to adopt a beta-like cell profile, and we next asked if transdifferentiation was associated with epigenetic changes. To this end, we analyzed DNA methylation in the human insulin gene region by bisulfite conversion assay. At three well-characterized DNA methylation sites in the human insulin locus ^29^ the +63, +127 and +139 positions DNA methylation was significantly reduced in ALDH3B2 mutant PANC-1 cells compared to control NTC-PANC-1 cells (Fig. 6a and 6b). For DNA methylation analysis of primary human pancreatic duct cells, we included primary human islets for comparison. Again, ALDH3B2 mutation significantly reduced DNA methylation at the same three sites in the insulin gene locus (Fig. 6c and 6d). However, ALDH3B2 mutation did not reduce the DNA methylation to the level observed in primary islets, a finding likely due to only a fraction (8-15%) of primary pancreatic duct cells were transdifferentiated into beta-like cells with ALDH3B2 mutation. Overall, DNA methylation analyses suggest that loss-of-function of ALDH3B2 caused epigenetic changes in the pancreatic duct cells to induce a stable cell fate change into pancreatic beta-like cells.

**Fig. 6.**
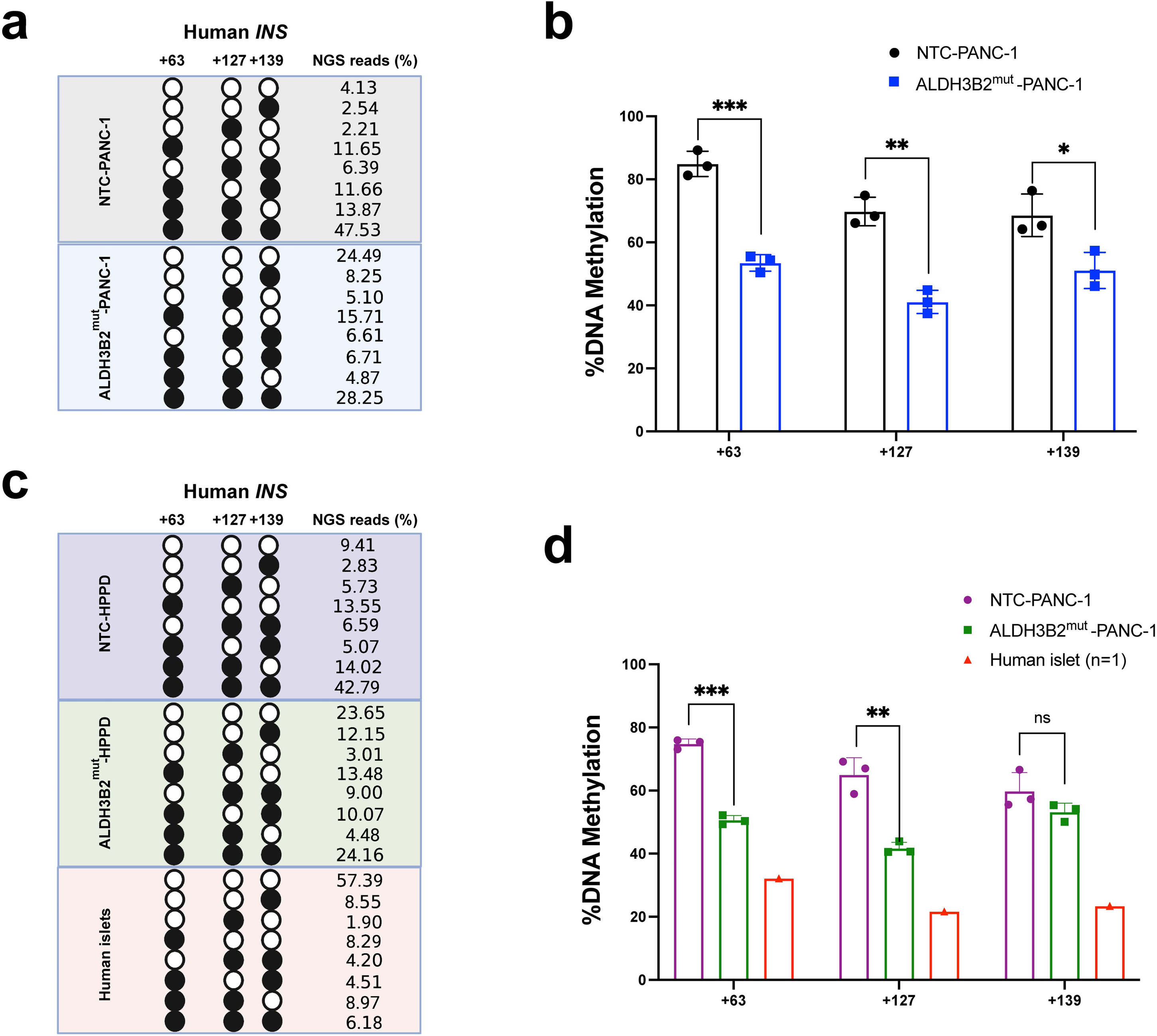
ALDH3B2 loss-of-function in pancreatic duct cells leads to epigenetic changes. **a,** Pattern and percentage of the DNA methylation at the position +63, +127 and +139 of the human insulin gene locus in NTC or ALDH3B2^mut^-PANC-1 cells. **b**, quantification of the DNA methylation percentage in **a**. **c**, Pattern and percentage of the DNA methylation at the position +63, +127 and +139 of the human insulin gene locus in HPPD-NTC, HPPD-ALDH3B2^mut^ and human primary pancreatic islet cells. **d**, quantification of the DNA methylation percentage in **c**.

### ALDH3B2 loss-of-function in human pancreatic duct cells induces heterogeneous beta-like cell populations with overlapping endocrine and duct cell identity

To investigate the characteristics of transdifferentiated beta-like cells in more detail, we performed 3’ gene expression single cell RNA sequencing of control or ALDH3B2 mutant HPPD cells and identified 13 unique cell clusters with a total of 3701 cells for the control and 4517 cells for the ALDH3B2 mutant condition analyzed (Fig. 7a). Whereas the control NTC-HPPD condition only showed a few (21 cells) insulin positive cells (relative expression intensity ≥ 1.0) most prominently in cluster 10, which could represent rare spontaneous transdifferentiated beta-like cells from duct cells, ALDH3B2 mutant HPPD cells develop insulin-expressing cells in various cell clusters, most prominently in clusters 5, 10, 11 and 12 with a total of 817 cells (Fig. 7b-d). The beta-like-cell-containing clusters 5, 10, 11 and 12 are a larger proportion of the ALDH3B2 mutant HPPD cells than in control NTD-HPPD cells (Extended Data Fig. 10a). The total percentage of insulin-expressing cells in ALDH3B2 mutant HPPD cells was 18.1 % but only 0.6 % in NTC HPPD cells; about 93% of all cells in both conditions still retained the expression of duct cell marker KRT19 (Extended Data Fig. 10b). The majority of insulin-positive cells also show significantly higher expression of other beta cell marker genes such as CHGA and IAPP but also duct cell identity marker genes KRT17, 19, and 23 (Fig. 7e). Differential gene expression analysis of insulin high-expressing cells (relative intensity >1) compared to insulin low-expressing cells (relative intensity <1) within the ALDH3B2 mutant HPPD cell condition showed significant upregulation of key beta cell marker genes such as CHGA, IAPP, SCGN, SCG3 and SCG5 (Fig. 7f). Of note, <u>g</u>ene <u>s</u>et <u>e</u>nrichment <u>a</u>nalysis confirmed the upregulation of key beta cell-specific gene sets related to peptide hormone metabolism, regulation of insulin secretion, and insulin processing (Fig. 7g). Re-analysis of insulin high-expressing cells in the ALDH3B2 mutant HPPD cells also identified heterogeneous cell populations with high beta cell identity (high INS/CHGA/IAPP co-expression), and polyhormonal cells (co-expression of INS/GCG/PPY), and endocrine progenitor-like cells (co-expression of INS and PAX6) (Extended Data Fig. 11).

**Fig. 7.**
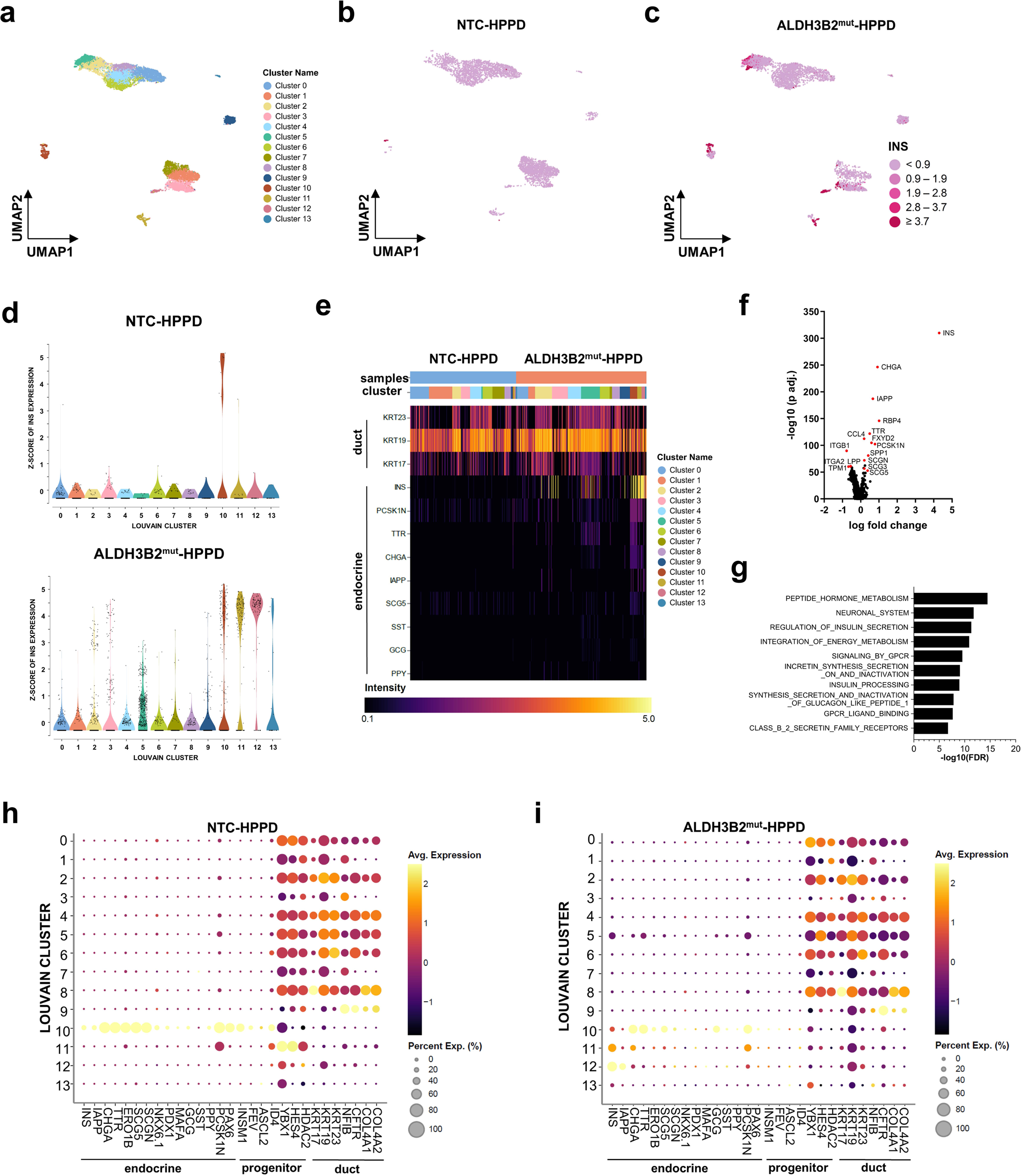
Single cell RNA sequencing of ALDH3B2 mutant human primary pancreatic duct cells identified multiple clusters of insulin producing beta-like cells with overlapping duct cell gene expression profiles. **a-c:** 13 unique cell clusters with a total of 3701 cells for the control and 4517 cells for the ALDH3B2 UMPA plots showing all identified cell cluster (**a**), insulin (INS) expression level of non-target control (NTC) (**b**) and ALDH3B2^mut^ (**c**) human primary pancreatic duct cells (HPPD). **d**, violin plot showing INS expression level of each individual cell grouped by cell cluster for NTC (left) and ALDH3B2^mut^ HPPD cells (right). In the ALDH3B2^mut^ 817 insulin-expressing cells were found, most prominently in clusters 5, 10, 11 and 12; in the NTC cells there were only 21. **e**, Heatmap showing indicated endocrine cell and duct cell maker gene expression of control (blue) and ALDH3B2^mut^ (red) HPPD cells at single cell level grouped by cell cluster. **f**, volcano plot showing differential expressed genes in ALDH3B2^mut^ HPPD cells comparing cells with high INS expression (intensity >1) vs. low INS expression (intensity <1). **g**, Gene set enrichment analysis showing the upregulated reactome of ALDH3B2^mut^ HPPD cells using the top 300 most significant upregulated genes of high INS-expressing (intensity >1) vs. low INS-expressing (intensity <1) cells. **h** and **i**, Dot plots showing average gene expression of indicated endocrine, progenitor, and duct cell marker genes in all identified cell cluster of NTC (**h**) or ALDH3B2^mut^ HPPD cells (**i**).

To further characterize the cell identity of insulin-expressing cells we compared the average gene expression of several marker genes specific for beta cells and other endocrine cells, endocrine progenitor cells, and duct cells ^30^ within all 13 cell clusters (Fig. 7h and 7i). In cluster 10 of the control condition, we detected insulin expression in about 40% of all cells (a total of 21 cells). The majority (60-80%) of the cells in this cluster have a low ductal cell-specific expression profile and high expression of endocrine progenitor cell markers, such as PAX6 and INSM1, suggesting that the majority of cells in cluster 10 may represent endocrine progenitor-like cells. Interestingly, ALDH3B2 loss-of-function in cluster 10 shifts cell identity towards beta-like cells, shown by downregulation of almost all endocrine progenitor marker genes and sustained expression of INS and CHGA. Cluster 11 and 12 show altered duct cell identity compared to cluster 0 to 9 even in the control condition and seem to have a higher percentage of beta-like cell transdifferentiation following loss-of-function of ALDH3B2. In comparison, cells in cluster 5 demonstrated strong duct cell identity but had high percentage of beta-like cell transdifferentiation as well.

To better understand why cells in cluster 5 and 12 have a higher percentage of transdifferentiation into beta-like cells, we performed trajectory inference analysis to identify the potential starting point of transdifferentiation (from low to high insulin expression, Extended Data Fig. 10c). Beta-like cells in cluster 5 may have originated from cluster 2 and cluster 5 itself. Re-analysis of cluster 5 revealed several insulin low-expressing cell cluster as possible starting points. However, insulin positive cells in cluster 5 show significant upregulation of genes involved in energetic processes, such as oxidative phosphorylation, aerobic respiration, and ATP synthesis, that may represent important prerequisites for beta-like cell transdifferentiation (Extended Data Fig. 10d and 10e). Cluster 3 may represent the originating cluster for beta-like cells in cluster 12. Differential gene expression analysis comparing cluster 2 and 3 (high potential for beta-like cell transdifferentiation) to cluster 4 (low potential of transdifferentiation) reveals that upregulation of genes important for translation and oxidative phosphorylation (cluster 2), enrichment of small GTPases RAC1/RHO (cluster 3), and elevated glycolysis (cluster 2 and 3) may favor beta-like cell transdifferentiation mediated by loss of function of ALDH3B2 (Extended Data Fig. 10f-k). In summary, ALDH3B2 loss-of-function may drive transdifferentation of pancreatic duct cells partially through duct cell-derived endocrine progenitor-like stage, and then into beta-like cells that still keep partial duct cell identity. Elevation of energy metabolism such as oxidative phosphorylation and glycolysis may be a key step for pancreatic duct cells to transdifferentiate into beta-like cells.

## DISCUSSION

In this study, we have described the first unbiased and genome-wide CRISPR screen in search for genes that regulate the transdifferentiation of human pancreatic duct cells into insulin-producing beta-like cells. We show that loss-of-function of a single gene, ALDH3B2, in human pancreatic duct cells is sufficient to drive them towards a beta-like cell fate. Although the pancreatic duct-to-beta cell transdifferentiation in human has been observed, as evidenced by the existence of INS^+^/CK19^+^ cells in pancreatic ductal epithelium, it is still a relatively rare event with the percentage of INS^+^ pancreatic duct cells estimated at ∼1% ^31^. Here we show that disruption of ALDH3B2 can drive transdifferentiation of primary human pancreatic duct cells into beta-like cells with an efficiency of ∼15%. This significant result carries potential for the development of a therapeutic intervention that could promote pancreatic duct-to-beta cell transdifferentiation for human beta cell mass replenishment.

We demonstrated that the trans-differentiated human pancreatic beta-like cells are functional, responsive to glucose challenge *in vivo*, and able to significantly lower blood glucose in diabetic animal models. Notably, neither ALDH3B2^mut^ PANC-1 cells nor HPPD cells were able to lower blood glucose to euglycemic levels in our studies. In comparison with primary human islets, the ALDH3B2 mutation transdifferentiated beta-like cells seem to have only modest expression of key beta cell markers (Fig. 4d-g), ability to secrete human insulin and lower blood glucose and in diabetic mice (Fig. 5a-c) and changes in DNA methylation status on insulin promoter (Fig. 6c and 6d). However, all these measurements and characterization were done on a mixed cell population with only about 15% of the cells being transdifferentiated beta-like cells. In theory, the actual changes in each transdifferentiated beta-like cell should be more dramatic than what our experimental data on the bulk cells showed. Our findings of immunofluorescent staining for insulin and key beta cell transcription factors and insulin granules with abundant ER ultrastructurally in the transdifferentiated beta-like cells strongly support the conclusion that loss-of-function of ALDH3B2 does induce *bona fide* cell transdifferentiation from human pancreatic duct cells into functional beta-like cells.

Similarly, it is likely that the quantity of transplanted transdifferentiated cells was insufficient to normalize blood glucose levels in these severely diabetic mice. Normally, approximately 2000 IEQ of human islets are required to fully normalize blood glucose levels in diabetic mice ^32^. In our experiments, we transplanted 10^7 HPPD cells per mouse. Considering that the ALDH3B2 gRNA lentiviral transduction efficiency in HPPD cells is about 40% and approximately 15% of the transduced cells eventually transdifferentiated into beta-like cells, we effectively transplanted around 600,000 beta-like cells per diabetic mouse. This is roughly equivalent to about 500 IEQ of human islets, given that each human IEQ contains an average of approximately 1200 beta cells ^33^. In both these diabetic mice and the control experiment in non-diabetic mice the transplanted ALDH3B2 ^mut^ HPPD cells had human insulin secretion and lowered the blood glucose values. We may be able to lower the blood glucose further in diabetic mice if a larger number of transdifferentiated cells were transplanted. To this end, we will need to improve lentiviral transduction efficiency and perhaps identify more effective gRNA sequences that target ALDH3B2 gene.

It is interesting that the ALDH family is often considered as a stem cell or progenitor marker. High aldehyde dehydrogenase activity (using Aldefluor assay ^34,35^) has been widely used to identify adult stem cells or progenitor cells in various tissues/organs, including hematopoietic stem cells (HSC) ^35^, neuronal progenitor cells (NPC) ^36,37^ or potential pancreatic progenitor cells ^38^. However, the Aldefluor assay cannot distinguish between the activity of different aldehyde dehydrogenases, so different ALDH members may be marking stem/progenitor cells in different tissues/organs. The function of ALDH3B2 is not yet well understood in any tissue. One rodent study reported that ALDH3B2 localizes to lipid droplets in cells and catalyzes the conversion of long-chain fatty aldehydes into long-chain fatty acids ^39^. How this function would impact pancreatic duct cell transdifferentiation into beta-like cells is unclear and warrants further studies.

The discovery of ALDH3B2 as a regulator of pancreatic duct-to-beta cell transdifferentiation provides a novel therapeutic target for pancreatic beta cell mass restoration. Endogenous pancreatic duct cells could potentially be targeted by gene-editing to mutate ALDH3B2 and induce transdifferentiation. Alternatively, ALDH3B2 enzymatic activity may be targeted by small molecules inhibitors to achieve the effects similar to those we observed using genetic disruption of the ALDH3B2 gene. The ALDH3B2 enzymatic assay established by Kitamura and colleagues ^39^ using long chain fatty aldehyde as substrate could be a useful chemical screen platform for potential new drug discovery. In support of this approach, the broad Aldehyde dehydrogenase (ALDH) inhibitors 4-diethylaminobenzaldehyde (DEAB) and Disulfiram (DSF) have been shown to promote beta cell differentiation in zebrafish and in PANC-1 cells ^40^, where it is possible that DEAB and DSF elicit their effect through inhibiting ALDH3B2. It will be of great importance to find out which aldehydes are unique substrates for ALDH3B2 and search for ALDH3B2-specific inhibitors, since mutation of the close member ALDH3A1 does not have the same effect (Fig. 1c and 1d) and another close member, ALDH1A3, was shown to be involved in the de-differentiation of pancreatic beta cells in Type 2 diabetes patients ^26^.

Our study identifying ALDH3B2 as a regulator of pancreatic duct-to-beta cell transdifferentiation was conducted using a human pancreatic duct cell line and primary human pancreatic duct cells. Given the difficulties of translating rodent studies into human in the beta cell regeneration research field, the findings presented here in human cells are directly relevant and have clear potential for the development of human diabetes therapeutics.

**Extended Data Fig. 1.**
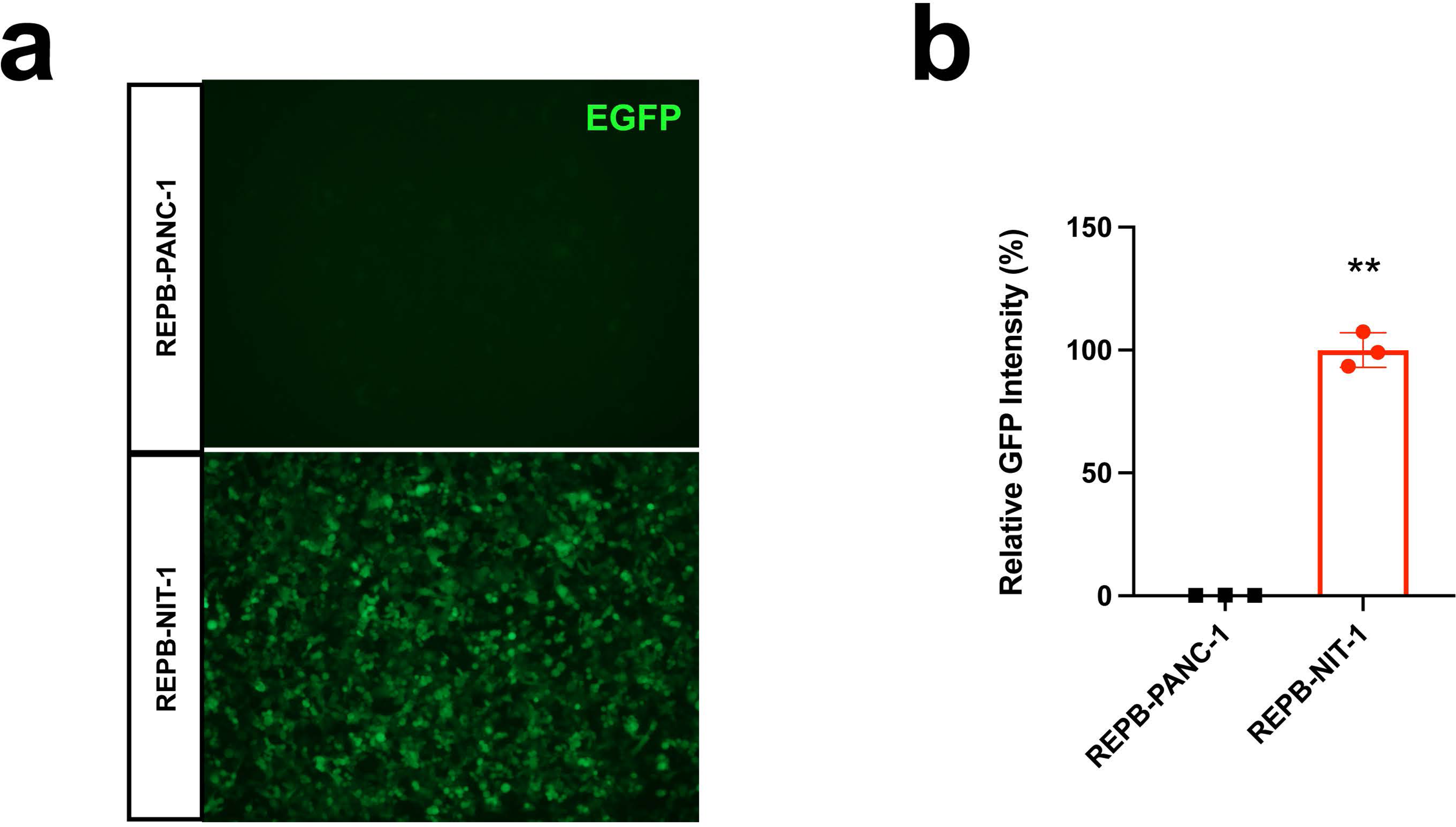
Characterization of the REPB reporter. EGFP expression by fluorescent imaging **(a)** and quantification of EGFP intensity **(b)** of the REPB reporter lentivirus infected PANC-1 and NIT-1 cells.

**Extended Data Fig. 2.**
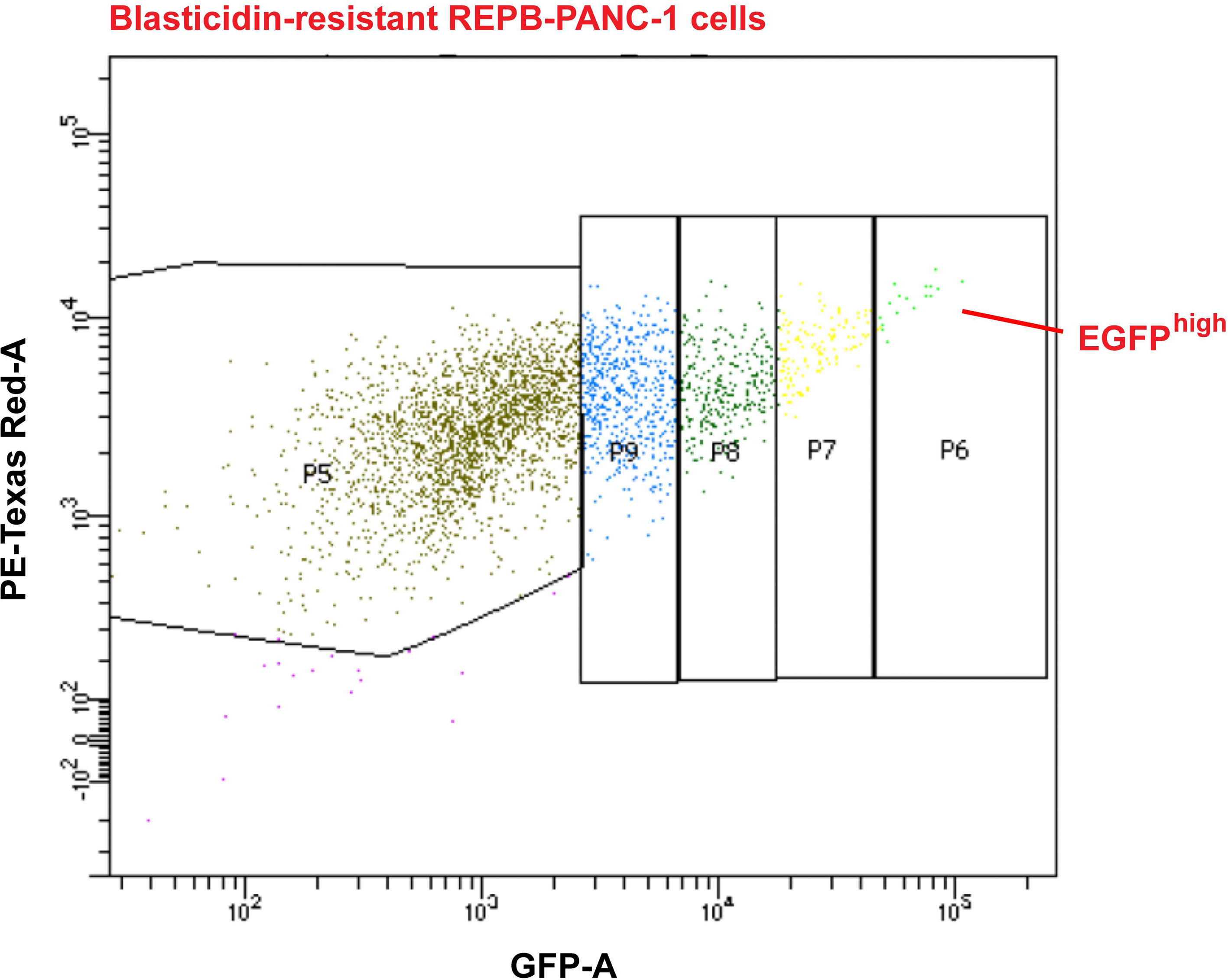
Flow cytometric sorting of the EGFP^high^ PANC-1 cells. Flow cytometry gating strategy of the genome-wide CRISPR screen. P6 population is considered as EGFP^high^ cells and isolated for subsequent sequencing.

**Extended Data Fig. 3.**
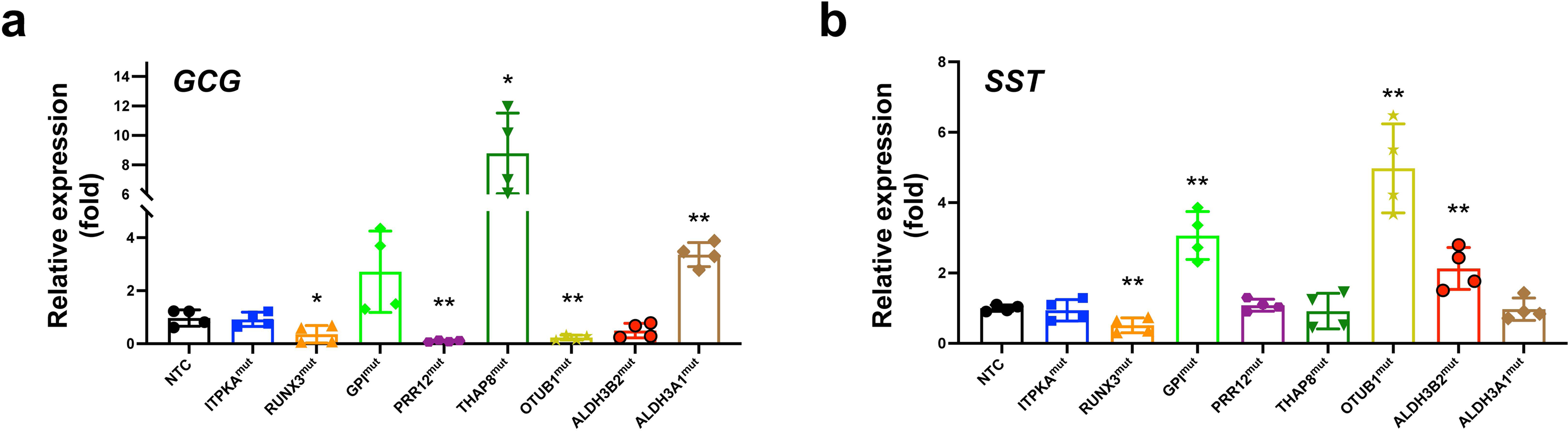
Quantification of GCG and SST expression. qPCR analysis of Glucagon (GCG) **(a)** and Somatostatin (SST) **(b)** expression level for top 8 candidate gene mutant PANC-1 cells.

**Extended Data Fig. 4.**
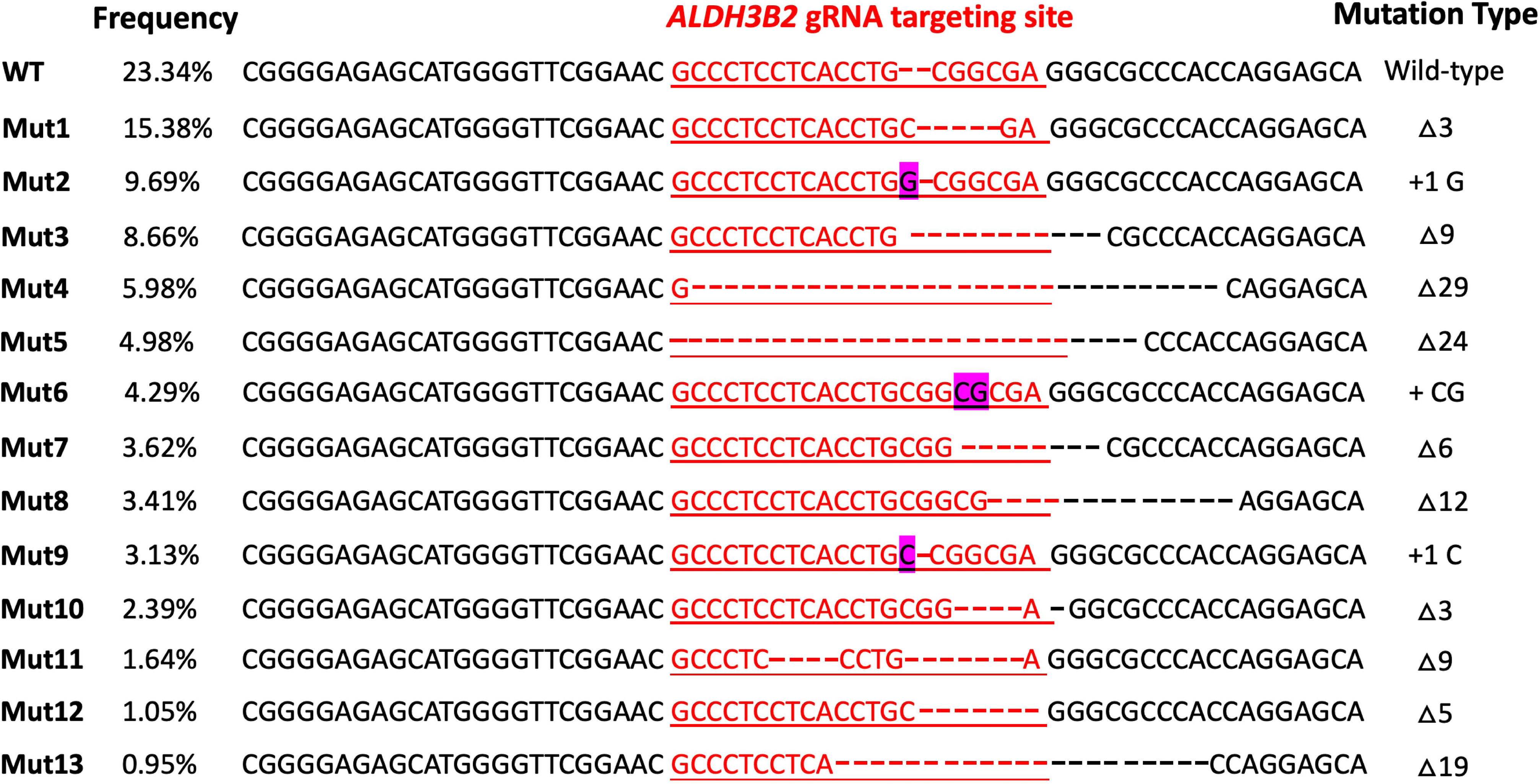
Indel analysis of the ALDH3B2^mut^-PANC-1 cells. The genome sequence flanking the gRNA targeting sites were sequenced and the most abundant indel mutations and their frequency is listed. The ALDH3B2 gRNA targeting site is labelled in red.

**Extended Data Fig. 5.**
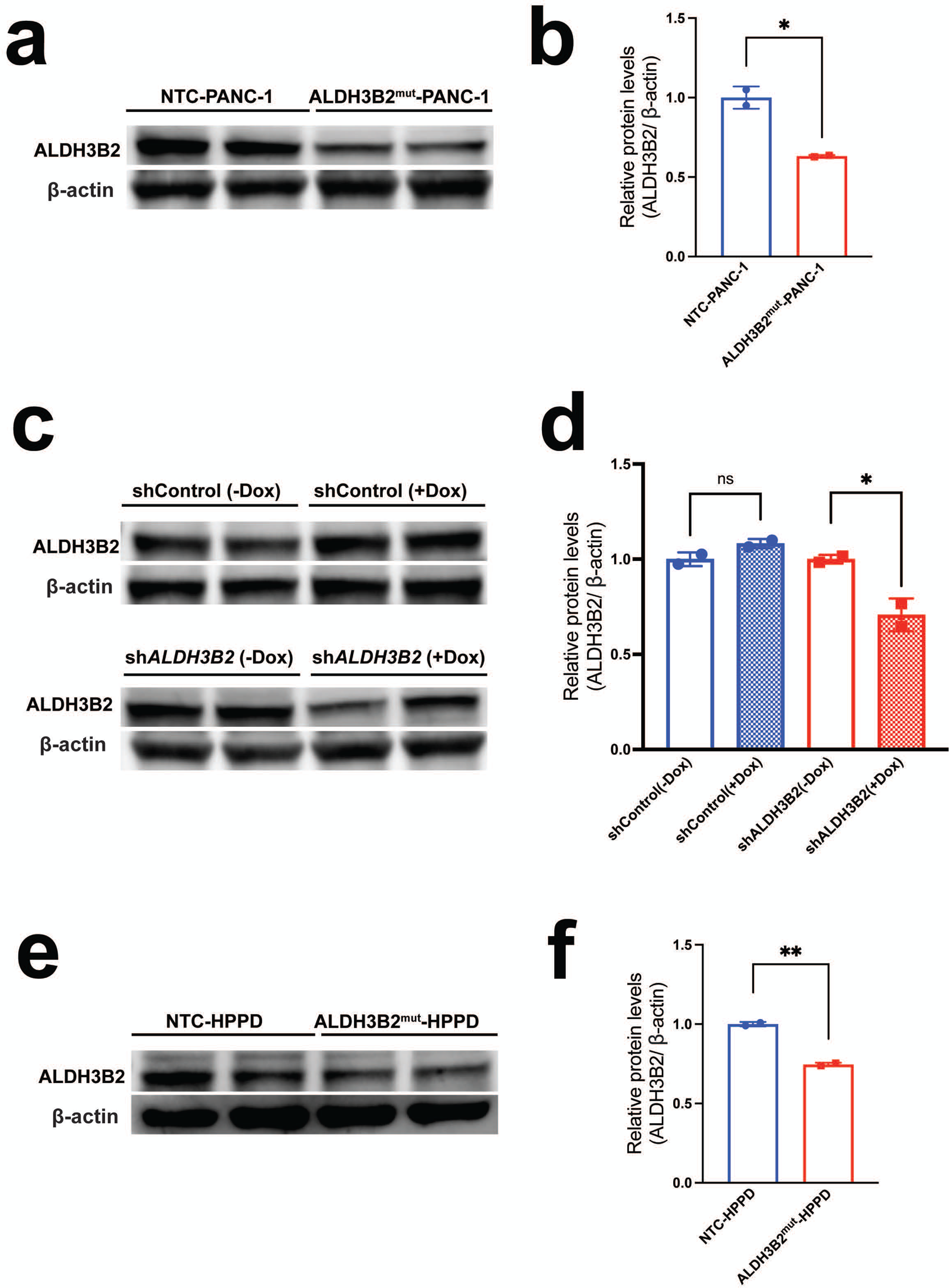
ALDH3B2 protein expression analysis in ALDH3B2 mutant or knockdown cells. **(a)**, Western blot shows that ALDH3B2 protein level is reduced in ALDH3B2 mutant PANC-1 cells compared to control NTC-PANC-1 cells. **(b)**, quantification of **(a)**. **(c),** Western blot shows that ALDH3B2 protein level is reduced only in PANC-1 cells with short hairpin RNA against ALDH3B2 (shALDH3B2) when treated with Doxycycline. **(d)**, quantification of **(c)**. **(e),** Western blot shows that ALDH3B2 protein level is reduced in ALDH3B2 mutant HPPD cells compared to control NTC-HPPD cells. **(b)**, quantification of **(a)**.

**Extended Data Fig. 6.**
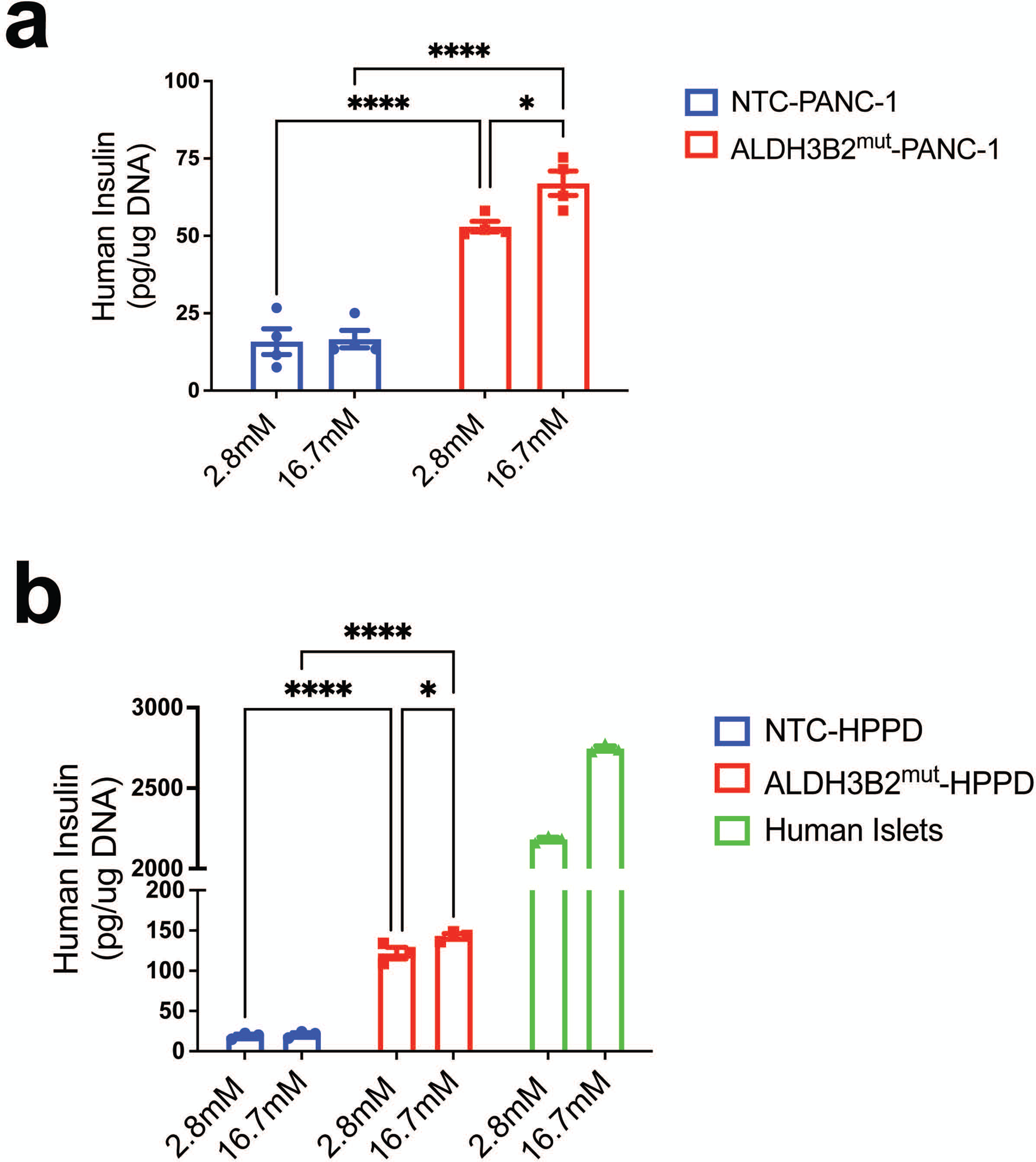
*in vitro* Glucose stimulated insulin secretion (GSIS) assay of ALDH3B2 mutant PANC-1 cells or HPPD cells. **(a)** Compared to control NTC-PANC-1 cells, the ALDH3B2 mutant PANC-1 cells has higher basal level insulin secretion (2.8mM), and increased insulin secretion upon high glucose treatment (16.7mM). Data show mean ± SEM of n=4. * p<0.05, ****p < 0.001. **(b)** ALDH3B2 mutant HPPD cells has higher basal level insulin secretion (2.8mM) than control NTC-HPPD cells, and mildly responds to high glucose treatment by increasing their insulin secretion (16.7mM). Human islets are included as a reference. Data show mean ± SEM of n=3. * p<0.05, ****p < 0.001.

**Extended Data Fig. 7.**
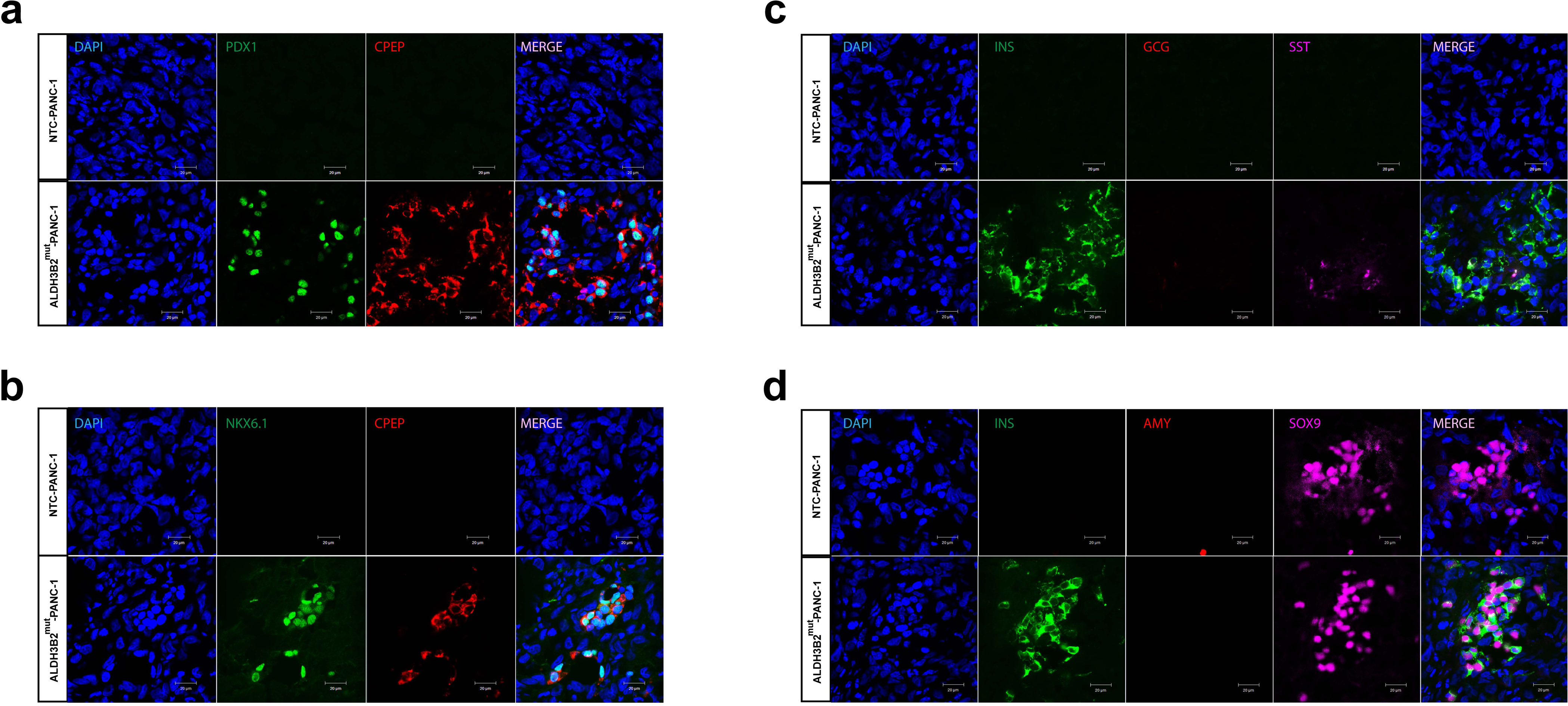
Immunofluorescence in transplanted NTC and ALDH3B2^mut^-PANC-1 cells. Pdx1 and C-Peptide immunofluorescence **(a)**, Nkx6.1 and C-Peptide immunofluorescence **(b)**, Insulin, Glucagon and Somatostatin immunofluorescence **(c)** and Insulin Amylase and Sox9 immunofluorescence **(d)** in transplanted NTC and ALDH3B2^mut^-PANC-1 cells.

**Extended Data Fig. 8.**
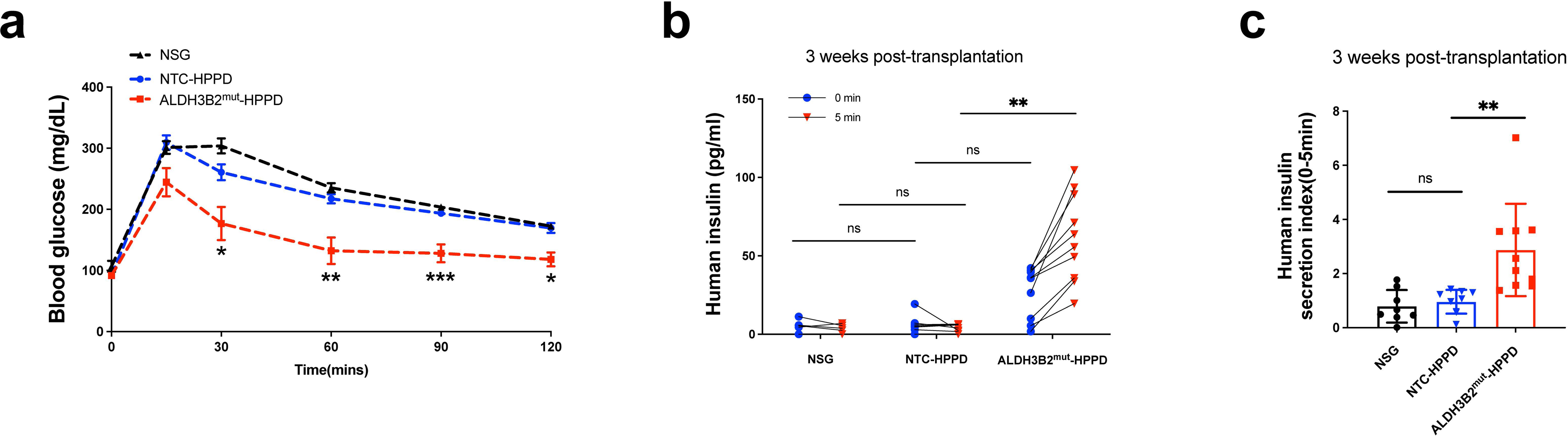
Functional characterization of NTC and ALDH3B2^mut^-HPPD cells transplanted in euglycemia NSG mice. **a,** An Intraperitoneal glucose tolerance test (IPGTT) in NTC or ALDH3B2^mut^-HPPD cells transplanted NSG mice 6 weeks post-transplantation. *p < 0.05, **p < 0.01, ***p < 0.001, calculated by two-way ANOVA with Sidak’s multiple comparisons test. **b,** Serum human insulin levels in non-transplanted NSG mice (Normal NSG), NTC-HPPD cells and ALDH3B2^mut^-HPPD cells transplanted NSG mice 5 minutes after intraperitoneal (IP) glucose injection at 3 week post-transplantation. **p < 0.01, calculated by two-way ANOVA with Sidak’s multiple comparisons test. **c,** Human insulin secretion index (0-5 minutes) calculated from **b**.

**Extended Data Fig. 9.**
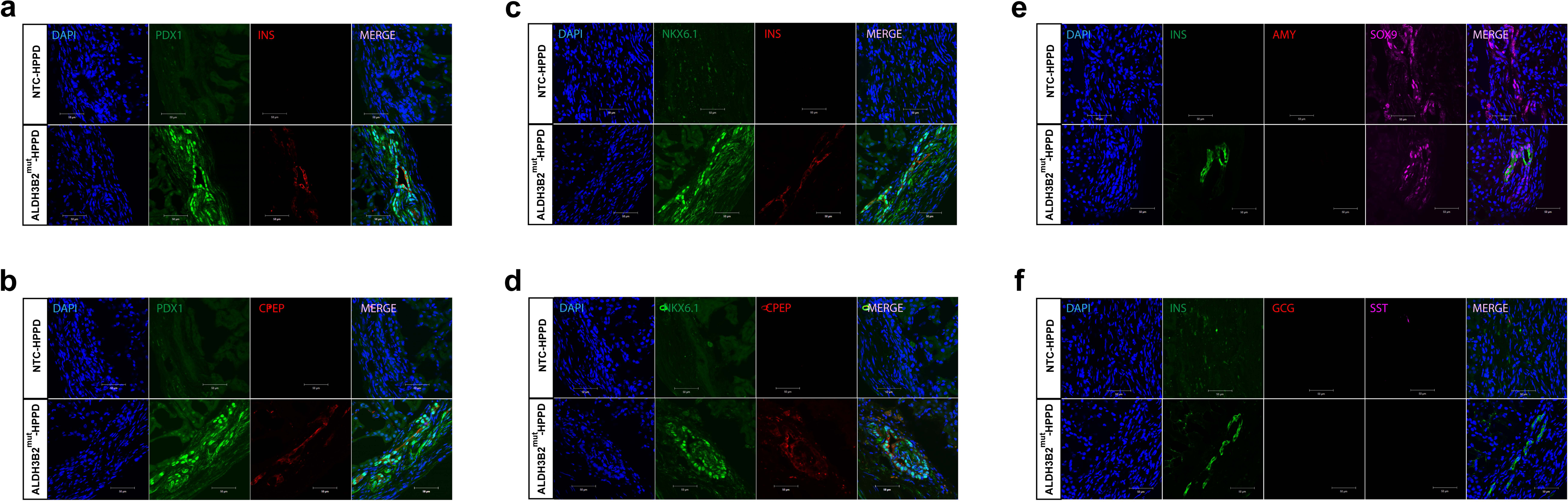
Immunofluorescence in transplanted NTC and ALDH3B2^mut^-HPPD cells. Pdx1 and Insulin immunofluorescence **(a)**, Pdx1 and C-Peptide immunofluorescence **(b)**, Nkx6.1 and insulin immunofluorescence **(c)**, Nkx6.1 and C-Peptide immunofluorescence **(d)**, Insulin Amylase and Sox9 immunofluorescence **(e)**, and Insulin, Glucagon and Somatostatin immunofluorescence **(f)** in transplanted NTC and ALDH3B2^mut^-HPPD cells.

**Extended Data Fig. 10.**
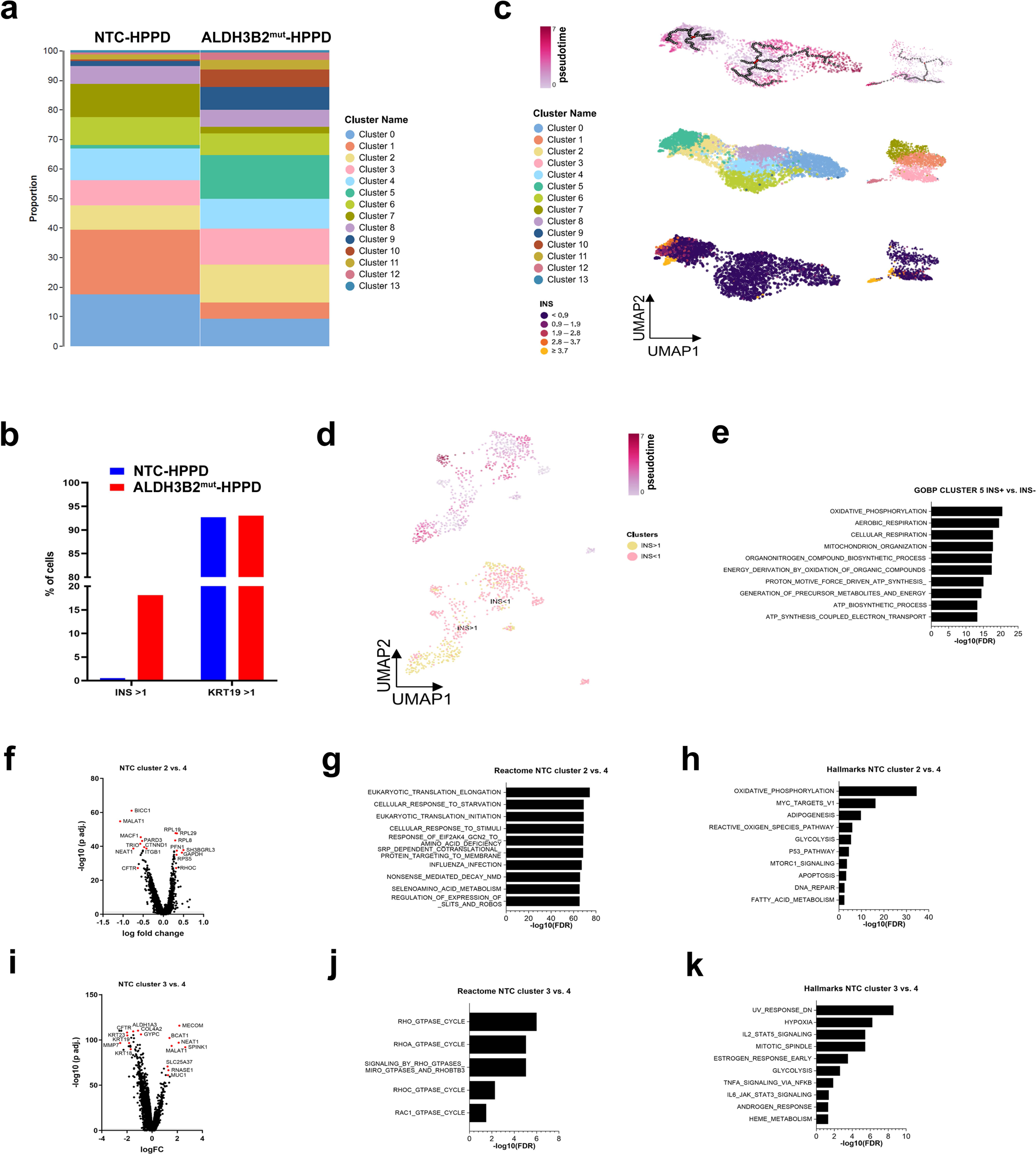
Cell cluster analysis from single cell RNA sequencing of ALDH3B2 mutant human primary pancreatic duct (HPPD) cells demonstrate multiple cellular origins and key characteristics of trans-differentiated HPPD-derived insulin (INS)-expressing beta-like cells. **a**, proportion of all identified cell cluster for non-target control (NTC, left) and ALDH3B2^mut^ (right) HPPD cells. UMAP plot showing all 13 identified cell cluster. **b**, proportion of INS and duct cell marker keratin 19 (KRT19) high (intensity >1) vs. low (intensity <1) expressing cells. **c**, Trajectory analysis showing calculated pseudotime (top) of indicated cell cluster (center) in comparison with insulin (INS) expression level (bottom). Black open circles represent calculated root nodes and red colored root nodes where defined as starting points. **d** and **e,** re-analysis of variable INS-expressing cluster 5 of ALDH3B2^mut^-HPPD cells showing UMAP plot of INS high (intensity >1, yellow dots) and INS low (intensity <1, pink dots) on the bottom and the trajectory analysis of the transition from INS low to INS high expressing cells in calculated pseudotime on the top (**d**). Gene ontology biological processes (GOBP) analysis of INS high vs. INS low ALDH3B2^mut^-HPPD cells from cluster 5 using the top 200 most significant upregulated genes (**e**). **f**-**k**, comparison of gene expression profiles and indicated pathway analysis between cluster 2 **(f-h)** and cluster 3 **(i-k)** showing high potential in beta-like cell transdifferentiation vs. cluster 4 showing no or low beta-like cell transdifferentiation capacity.

**Extended Data Fig. 11.**
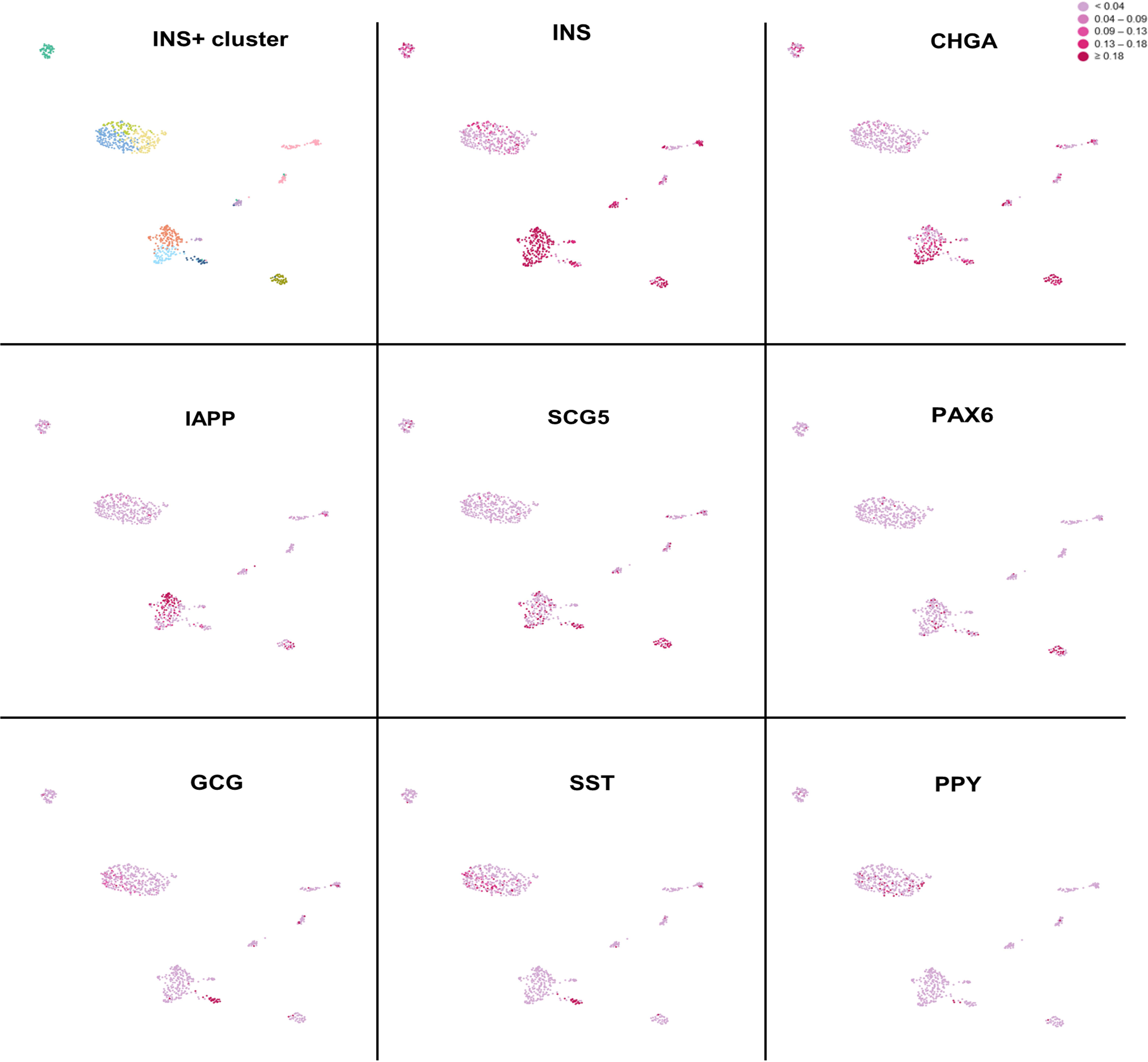
Key gene expression profiles of Inulin positive (intensity >1) cells from single cell RNA sequencing of ALDH3B2 mutant human primary pancreatic duct (HPPD) cells at single cell level. Cell cluster identification (top left) and gene expression levels of indicated endocrine marker genes (rest). Insulin (INS), Chromogranin A (CHGA), Islet Amyloid Polypeptide (IAPP), Secretogranin V (SCG5), Paired box protein Pax-6 (PAX6), Glucagon (GCG), Somatostatin (SST), Pancreatic polypeptide (PPY).

**Extended Data Fig. 12.**
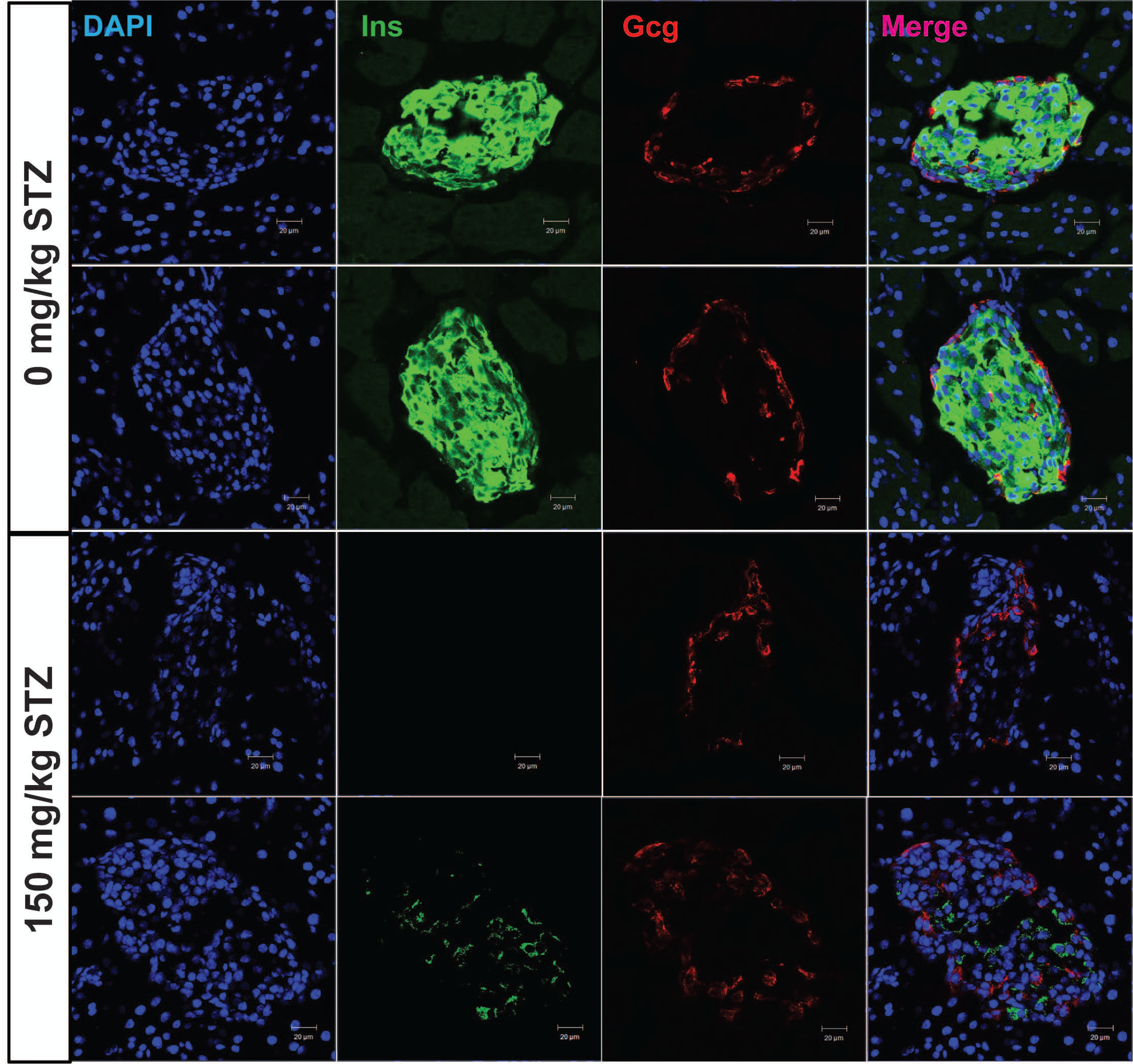
Beta cell specific destruction by high dose streptozotocin (STZ) in NSG mice. Immunofluorescence shows that 150mg/kg STZ injection in NSG mice leads to beta cell specific destruction (Ins immunostaining), while alpha cells remains unaffected (Gcg immunostaining).

## MATERIALS AND METHODS

### Mice

NSG (NOD.Cg-Prkdc^scid^ Il2rg^tm1Wjl^/SzJ) mice were purchased from the Jackson Laboratory (Bar Harbor, ME). Animals were housed in pathogen-free facilities at the Joslin Diabetes Center and all experimental procedures were approved and performed in accordance with institutional guidelines and regulations.

### REPB reporter construction and REPB PANC-1 cell generation

The REPB reporter lentivirus vector was constructed by assembling Rat insulin promoter, RIP3.1 promoter ^22^, EGFP and Blasticidin-S deaminase (BSD). The EGFP and BSD genes are fused together with P2A peptide. The REPB reporter lentivirus was used to infect PANC-1 cells (ATCC #CRL-1469), and the infected PANC-1 cells are then single-cell sorted by FACS. PANC-1 clones with confirmed REPB reporter genome incorporation were used in the genome-wide CRISPR screen.

### CRISPR GeCKO library screen

The human GeCKO-v2 (Genome-Scale CRISPR Knock-Out) lentiviral pooled library was obtained from Addgene (Addgene, # 1000000048) and was prepared as previously described ^41^. 100 million REPB PANC-1 cells were infected with human GeCKO CRISPR lentiviral library at MOI of 0.3, and then subsequently selected with puromycin (2.5 μg/ml) at day 3 post lentiviral infection. After cells recover from puromycin selection, 20 million cells were collected as baseline control (CON-1 and CON-2), the rest of the cells were further selected with blasticidin (10 μg/ml) for 7 days. The blasticidin-resistant cells were allowed to grow back to full confluence and then subjected to FACS sorting on their EGFP intensity. Cell population with the highest EGFP intensity were collected as experiment group (EXP-1 and EXP-2). Genomic DNA was extracted from the cells (Quick-gDNA midiprep kit, Zymo Research), the NGS (Next Generation Sequencing) libraries were prepared as previously described ^42^, and then subjected to NGS sequencing analysis (Novogene, CA). The gRNA sequences from the NGS sequencing data were extracted using standard bioinformatics methods, and the read count of gRNAs were calculated as Count Per Million (CPM).

### Cell lines

PANC-11 (#CRL-1469) and 293FT (#R7007) cell lines were obtained from ATCC and Thermo Fisher Scientific, respectively. Cells were maintained in DMEM (Gibco, 10313039), supplemented with 10% fetal bovine serum (FBS, Gibco), L-alanyl-L-glutamine (Gibco) and penicillin/streptomycin (Corning), in a 37° C incubator with 5% CO_2_. To generate non-targeting control (NTC) and ALDH3B2^mut^ PANC-1 cells, non-targeting (NTC) gRNA (5’-GCTTTCACGGAGGTTCGACG-3’) or ALDH3B2 gRNA (5’-GCCCTCCTCACCTGCGGCGA-3’, HGLibA_01571) oligos were cloned into LentiCRISPR-v2 vector. Wild type PANC-1 cells were then transduced by NTC or ALDH3B2 gRNA-containing lentivirus, and subsequently selected by puromycin treatment. Indel mutation in ALDH3B2^mut^ cells was confirmed by deep sequencing analysis (MGH DNA Core Facility, Cambridge, MA). All plasmid sequences were verified by Sanger sequencing before transduction and transfection. To generate shControl and shALDH3B2 PANC-1 cells, a scrambled or ALDH3B2-targeting shRNA was cloned into FH1t(INSR)UTG-GFP vector (A gift from Dr. Stephan Kissler). shControl and shALDH3B2 lentiviruses were then used to infect PANC-1 cells and the cells were selected by puromycin treatment.

### PANC-1 cells transplantation studies

Experimental diabetes was induced in 8-week-old NSG male mice by a single intraperitoneal injection of streptozotocin (STZ) (150 mg STZ/kg body weight) which resulted in the majority of the beta cells being killed (Extended Data Fig. 12). Animals were considered diabetic only if morning-fed blood glucose exceeded 350 mg/dl. Three days after the STZ injection, ∼10^7^ PANC-1 cells (carry NTC and ALDH3B2^mut^) were transplanted subcutaneously into each diabetic NSG mouse. Blood glucose was monitored every 3-4 days. Six weeks post cells transplantation, Intraperitoneal glucose tolerance test (IPGTT) was performed. Mice were fasted for 16 hours. The plasma glucose levels of the mice before (baseline) or 15, 30, 60, 90, and 120 minutes after intraperitoneal injection of 2 mg/g body weight glucose were recorded by a Glucose Meter. At day 52 post cells transplantation, grafts were surgically removed from NSG mice, and the blood glucose was measured 4 days later.

### Human pancreatic ductal cell isolation and purification

Human primary pancreatic ductal cells isolation was performed as previously described ^27,43^. In brief, “Human acinar”(which is actually islet depleted exocrine tissue) from Integrated Islet Distribution Program (IIDP) (Donor information is summarized in Supplemental Table 2) was washed 2 times with PBS, and then incubated with Trypsin solution (1.5ml 0.25% Trypsin in 20ml PBS) shaking at 37°C for 15 min. Dispersed cells were centrifuged at 1,000 rpm for 5 min, the supernatant was aspirated, and then the pellets were resuspended with mouse antihuman CA19-9 antibody (Invitrogen; clone 116-NS-19-9) in 2 ml PBS solution. After 15 min incubation at 4°C, the cell suspension was mixed with 10ml PBS solution (375mg EDTA and 2.5g BSA in 500ml PBS) gently. Tubes were centrifuged at 1,000 rpm for 5 min, supernatant was aspirated, 250 μl/tube goat anti mouse IgG microbeads (Miltenyi Biotec) in PBS solution were added, and pellets were mixed. After 20 min incubation at 4°C, wash with PBS solution 2 times. Tubes were centrifuged at 1,000 rpm for 5 min, and pellets were resuspended in 20 ml cold PBS solution and passed through 40 m cell strainers to remove newly formed clumps of cells. MACS magnetic LS separation columns (Miltenyi Biotec) were prepared according to the manufacturer’s instructions.

### Preparation and transplantation of primary human pancreatic duct cells

Human primary pancreatic ductal cells isolation was performed as above described. Purified human primary pancreatic ductal cells (HPPD) were immediately cultured in a low-attachment plate in RMPI DMEM/F12 medium (Gibco), supplemented with 10% FBS and penicillin/streptomycin. Lentivirus encoding a NTC or ALDH3B2 gRNA together with Cas9 endonuclease was added to the culture media for overnight infection. The next day, HPPD cells were washed with culture media twice and ∼10^7^ cells were transplanted under the left kidney capsule of 8-week-old of STZ induced diabetic male NSG mice. Graft recipients were left to recover from surgery for three weeks. At day 56 post-transplantation, the kidney transplanted with grafts were surgically removed for gene expression analysis by immunofluorescence. The final blood glucose measurement of NSG mice was done 4 days later.

### Western Blotting

Cellular lysates were gathered on ice employing RIPA lysis buffer enriched with proteinase and phosphatase inhibitors (Sigma-Aldrich, #11836153001). The concentration of proteins was determined utilizing the Pierce BCA protein assay (Thermo Scientific Fisher, #23225). Thirty micrograms of denatured cell lysate proteins were loaded for SDS-PAGE electrophoresis (4–20% TGX gel, Bio-Rad). Primary antibodies include ALDH3B2 (Proteintech, #15746-1-AP) and β-Actin (8H10D10) Mouse mAb (HRP Conjugate) (Cell signaling, #12262S). Peroxidase (HRP) Anti-Rabbit IgG Goat Secondary Antibody was used. A C-DiGit blot scanner and Image Studio software (LI-COR Biosciences) were performed for imaging and quantification.

### Glucose-stimulated insulin secretion (GSIS)

Cells or islets were washed twice with Krebs Ringer Bicarbonate HEPES (KRB) buffer containing 2.8 mM glucose, then incubated at 37°C in a 2.8 mM glucose KRB buffer for 1 hour. The supernatant was collected and saved for insulin measurement. The rest of the cells or islets were replaced with 16.8 mM glucose KRB buffer for another 1-hour incubation at 37°C. Collecting supernatant for insulin measurement, then cells or islets were incubated with 30 mM KCl and 16.8 mM glucose for 1 hour at 37°C before the final insulin sampling. Genomic DNA was purified from cells or islets to normalize insulin levels to DNA content. Insulin levels were measured by STELLUX® Chemi Human Insulin ELISA kit (Alpco, #80-INSHU-CH01).

### Quantitative real-time PCR (qPCR)

Cells or grafts were treated with TRIzol (Thermo Fisher Scientific) for RNA extraction following the manufacturer’s protocol. Purified RNA was reverse-transcribed into cDNA using the SuperScript IV first-strand synthesis kit (Invitrogen). INS (Hs00355773_m1), GCG (Hs01031536_m1), SST (Hs00356144_m1) PDX1(Hs00236830_m1), NKX6-1 (Hs00232355_m1), GCK (Hs01564555_m1), SLC2A2 (Hs00165775_m1), SLC2A1 (Hs00892681_m1), CPE (Hs00960598_m1), CA2 (Hs01070108_m1), KRT19 (Hs00761767_s1), SOX9 (Hs00165814_m1), ALDH3B2 (Hs02511514_s1) and ACTB (Hs01060665_g1) probes for TaqMan assays were purchased from Thermo Fisher Scientific. All Gene expression levels were analyzed by SYBR green PowerUp qPCR assays (Applied Biosystems). Primer sequences used is shown in the following table:

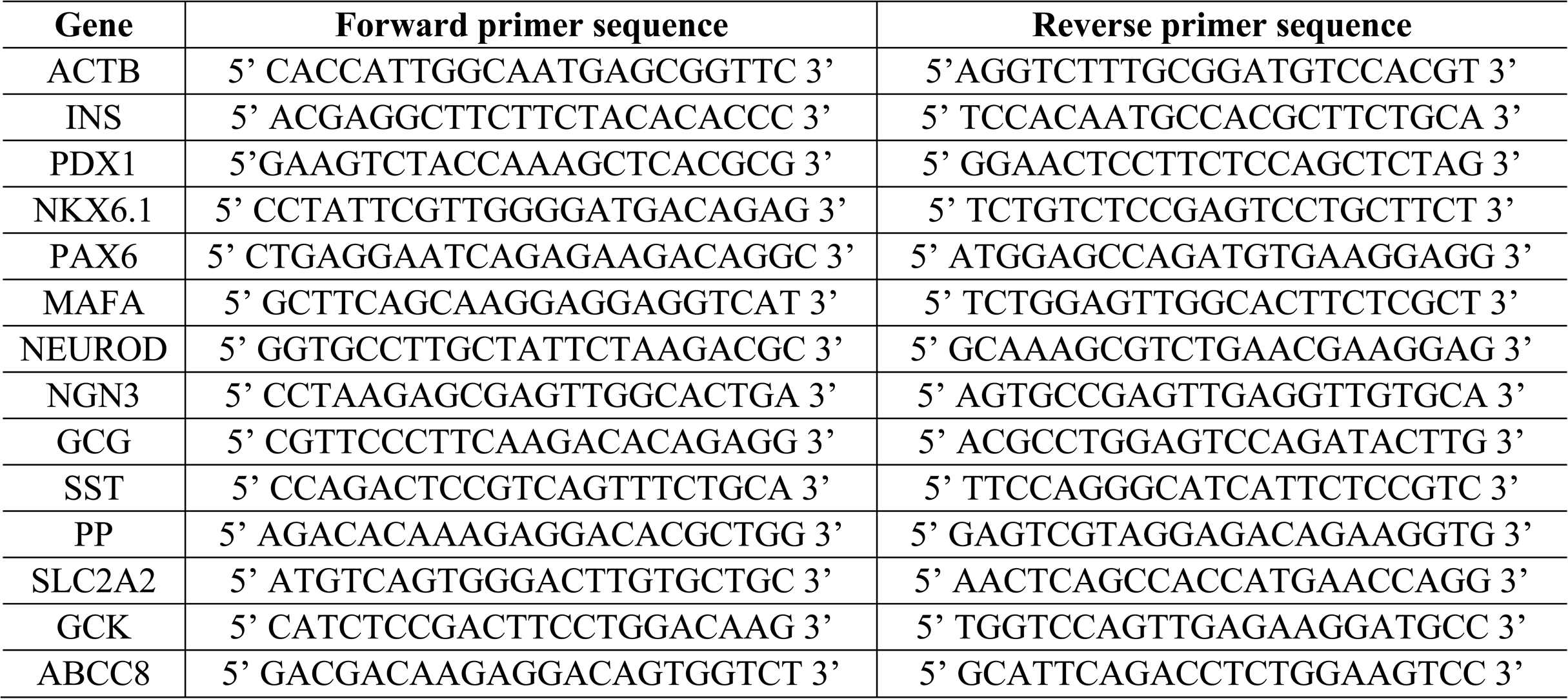
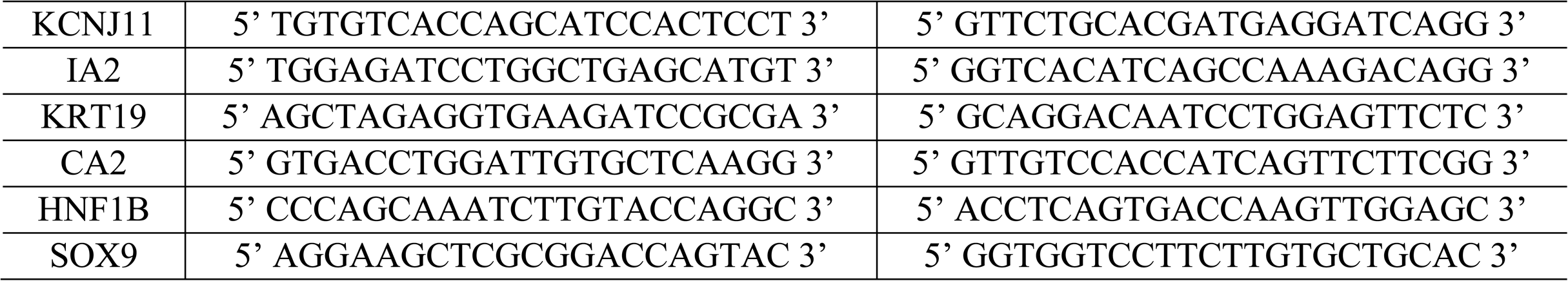

All qPCR assays were performed using a QuantStudio 6 Flex Real-Time PCR system (Applied Biosystems).

### Immunofluorescence staining and confocal microscopy

The subcutaneously transplanted PANC-1 cells or the kidney with HPPD graft transplantation were surgically removed from the mice, fixed 1 hour in 4% paraformaldehyde at 4°C, and dehydrated using 30% sucrose solution overnight. The tissues were embedded in disposable base molds (Thermo Fisher Scientific) and 10 mm sections were cut. For staining, slides were blocked with PBS+0.1% Triton X-100 (Thermo Fisher Scientific) +5% donkey serum (Sigma-Aldrich) for 1 hr at room temperature (RT), incubated with primary antibodies overnight at 4°C, washed, incubated with secondary antibody incubation for 1 hr at RT, incubated with Hoechst 33342 (Invitrogen) for 10 min at RT, and washed. For imaging, samples were mounted in fluorescence mounting medium (Dako), covered with coverslips, and sealed with nail polish. Representative images were taken using a Zeiss LSM 710 confocal microscope. Primary antibody: Insulin (A0564, Dako), Pdx1 (5679S, Cell Signaling Technology), Dykddddk Tag (14793S, Cell Signaling Technology), C-peptide (GN-ID4, Developmental Studies Hybridoma Bank) and Cytokeratin 19 (Abcam, ab7754). Cells were seeded into 4-well culture slide (Falcon) at density of 10^5^ cells/well. After another 24 hours, the cells were fixed, stained and subjected to fluorescence microscopic analysis as above described.

### Serum insulin measurement

Mouse blood was collected from the tail tip and allowed to clot. Serum was separated by brief centrifugation according to standard protocol. The serum insulin level was measured using STELLUX® Chemi Human Insulin ELISA kits (Alpco);the detection limit is 20pg/ml.

### Electron microscopy

NTC-PANC-1 or ALDH3B2^mut^-PANC-1 were fixed at RT for 2 hr with a mixture containing 1.25% PFA, 2.5% glutaraldehyde, and 0.03% picric acid in 0.1M sodium cacodylate buffer (pH 7.4). Samples were then sent to the Advanced Microscopy Core of Joslin for further processing and transmission electron microscope imaging.

### DNA methylation analyses

Bisulfite conversion (Zymo Research, EZ DNA Methylation-Direct Kits) of DNA from PANC-1, human primary pancreatic ductal cells carry NTC or ALDH3B2^mut^ and human islets were performed as described previously ^44^. Bisulfite-treated DNA was PCR amplified, using primers (human INS promoter forward primer - 5’ AGGATAGGTTGTATTAGAAGAGGTTATTAAG 3’; human INS promoter reverse primer- 5’ CCCCTAAACTCACCCCCACATACTTC 3’) specific for bisulfite treated DNA but independent of methylation status at monitored CpG sites. Reaction conditions for the first round of PCR were 5 cycles of 95 °C 1 min, 52 °C 3 min, 72 °C 3 min followed by 40 cycles of 95 °C 30 s, 55 °C 45 s, 72 °C 45 s followed by 7 min at 72 °C. PCR products were gel purified and used for deep sequencing analysis (MGH DNA Core Facility, Cambridge, MA).

### Single cell RNA sequencing

Isolation of human primary pancreatic ductal cells was performed as described above. Purified human primary pancreatic ductal cells (HPPD) were immediately cultured in a low-attachment plate in RMPI DMEM/F12 medium (Gibco), supplemented with 10% FBS and penicillin/streptomycin. Lentivirus encoding a NTC or *ALDH3B2* gRNA together with a Cas9-mCherry reporter construct was added to the culture media. 6 days later cells were harvested and live mCherry positive cells were isolated by FACS sorting. The sorted cells were used for scRNAseq using the Chromium Next GEM Single Cell 3’ GEM, Library & Gel Bead Kit v3.1 (cat # PN-1000213; 10 x Genomics) according to manufactures’ instructions. Illumina NovaSeq 6000 with about 1.3 billion reads total was used for sequencing the purified 3’ gene expression library. The single cell RNA-seq dataset was processed, explored and visualized using Cellenics® community instance (https://scp.biomage.net/) that’s hosted by Biomage (https://biomage.net/) using preset standard filtering thresholds and integration methods. Pre-filtered count matrices were uploaded to Cellenics. Dead or dying cells were removed by filtering droplets with high mitochondrial content (3% cut-off). Outliers in the distribution of number of genes vs number of UMIs were removed by fitting a linear regression model (p-values between 1.97E-04 – 2.34E-04). Cells with a high probability of being doublets were filtered out using the scDblFinder method (threshold range: 0.50 - 0.56). Overall filtering rates after processing are in the range of 10.75% - 11.35% of cells. Data normalization, principal-component analysis (PCA) and data integration using Harmony were performed on data from high-quality cells. Clusters were identified using the Louvain method, and a Uniform Manifold Approximation and Projection (UMAP) embedding was calculated to visualize the results. Cluster-specific marker genes were identified by comparing cells of each cluster to all other cells using the presto package implementation of the Wilcoxon rank-sum test.

### Statistical analyses

Statistical analyses were performed by unpaired or paired tests as indicated using the Prism 8 software. All data are presented as mean ± SEM. Significance was defined as *p < 0.05, **p < 0.01 and ***p < 0.001. No samples were excluded from the analysis. Data analysis was not blinded. All data are representative of two or more similar experiments.

## ACKNOWLEDGEMENTS

This research was supported by funds from the Beatson Foundation to P.Y and S.B-W, by postdoctoral fellowship to J. L. from the Mary K. Iacocca Foundation and JDRF, and the Diabetes Research and Wellness Foundation Chair to S.B-W. We acknowledge the support from core facilities funded both by the NIDDK Diabetes Research Center (P30DK036836) and the Joslin Diabetes Center, especially the Gene editing and CRISPR screen core facility. We acknowledge the generous comments and editing from Dr. Gordon Weir and Dr. Stephan Kissler of Joslin Diabetes Center. We also acknowledge Wuhan University for supporting J. L.’s research, which contributes to the completion of his Ph.D. at the Key Laboratory of Combinatorial Biosynthesis and Drug Discovery, School of Pharmaceutical Sciences, Wuhan University, China.

