## Supplemental Table 2 for "Loss-of-function of ALDH3B2 transdifferentiates human pancreatic duct cells into beta-like cells"

### IIDP Research Data Repository Custom Data Report

| RRID | Tissue types | Donor Information | | | | | | Study Information | | | Experiment design | | | | | | | |
|---|---|---|---|---|---|---|---|---|---|---|---|---|---|---|---|---|---|---|
| | | Demographics | | | Clinical Information | | | | | | | | | | | | | |
| | | Sex | Age at death (Yrs) | Race/Ethnicity | Height (cm) | Weight (kg) | BMI | Offer Type | Shipment Date (UTC) | Isolation ID | qPCR | IF staining | Western Blot | In vivo Tx | GSIS | DNA methylation sequencing | Electron Microscopy | Single-cell RNA-seq |
| RRID:SAMN13023422 | Exocrine | Female | 57.0 | White | 169.0 | 67.0 | 23.5 | Fresh | 10/14/19 6:00 PM | HU1180 | O | | | | | | | |
| RRID:SAMN13841476 | Exocrine | Female | 30.0 | Hispanic/Latino | 163.0 | 72.0 | 27.1 | Fresh | 1/14/20 8:00 PM | HP2327 | | O | | | O | | | |
| RRID:SAMN16705048 | Exocrine | Male | 24.0 | Hispanic/Latino | 180.0 | 78.8 | 24.3 | Fresh | 11/9/20 4:00 PM | HU1208 | O | | | O | | | | |
| RRID:SAMN17277414 | Exocrine | Female | 43.0 | Hispanic/Latino | 156.0 | 88.8 | 36.5 | Fresh | 1/11/21 5:00 PM | HU1212 | | O | | | | O | | |
| RRID:SAMN19588420 | Exocrine | Male | 61.0 | Hispanic/Latino | 162.0 | 77.0 | 29.3 | Fresh | 6/7/21 4:00 PM | HU1221 | O | | O | | | | | |
| RRID:SAMN20926065 | Exocrine | Female | 62.0 | Black or African American | 158.0 | 67.0 | 26.8 | Fresh | 8/23/21 5:00 PM | HU1224 | | | | O | | O | | |
| RRID:SAMN22814514 | Exocrine | Male | 26.0 | White | 192.0 | 107.5 | 29.2 | Fresh | 11/1/21 4:00 PM | HU1229 | | O | | | | O | | |
| RRID:SAMN27361473 | Exocrine | Male | 34.0 | Hispanic/Latino | 177.0 | 99.7 | 31.8 | Fresh | 4/6/22 4:00 PM | HU1238 | | | | O | | | | |
| RRID:SAMN28867623 | Exocrine | Male | 36.0 | White | 179.0 | 94.8 | 29.6 | Fresh | 6/6/22 4:00 PM | HU1244 | O | O | | | | | | |
| RRID:SAMN32641506 | Exocrine | Male | 16.0 | Hispanic/Latino | 185.0 | 101.0 | 29.5 | Fresh | 1/10/23 1:00 AM | HU1256 | | | O | | O | | O | |
| RRID:SAMN32869720 | Exocrine | Female | 29.0 | Hispanic/Latino | 165.1 | 87.0 | 32.0 | Frozen | 1/23/23 6:00 AM | UWHI330R | | | | | | | | |
| RRID:SAMN33397516 | Exocrine | Male | 63.0 | White | 172.7 | 65.8 | 22.0 | Fresh | 2/21/23 10:30 PM | HP-23052-01 | | | | | | | | O |
| RRID:SAMN37567961 | Exocrine | Male | 22.0 | Hispanic/Latino | 177.0 | 64.0 | 20.4 | Fresh | 9/27/23 7:00 PM | HU1276 | | | | | | | | O |
| RRID:SAMN20478103 | Islet | Male | 30.0 | Hispanic/Latino | 182.9 | 84.9 | 25.4 | Fresh | 7/29/21 9:45 PM | N/A | | | | | | O | | |
| RRID:SAMN32683021 | Islet | Female | 20.0 | Hispanic/Latino | 155.0 | 87.5 | 36.4 | Fresh | 1/12/23 6:00 PM | N/A | O | | | | O | | O | |
